# Dopamine and serotonin differentially associated with reward and punishment processes in humans: A systematic review and meta-analysis

**DOI:** 10.1101/2025.01.08.631868

**Authors:** Anahit Mkrtchian, Zeguo Qiu, Yaniv Abir, Tore Erdmann, Quentin Dercon, Terezie Sedlinska, Michael Browning, Harry Costello, Quentin J. M. Huys

**Author notes:** Corresponding Author, Anahit Mkrtchian, UCL Division of Psychiatry, 149 Tottenham Ct Rd, London, W1T 7NF, UK.

## Abstract

**Importance:** To support treatment assignment, mechanistic biomarkers should be selectively sensitive to specific interventions. Here, we examine whether different components of reinforcement learning in humans satisfy this necessary precondition. We focus on pharmacological manipulations of dopamine and serotonin that form the backbone of first-line management of common mental illnesses such as depression and anxiety.

**Objective:** To perform a meta-analysis of pharmacological manipulations of dopamine and serotonin and examine whether they show distinct associations with reinforcement learning components in humans.

**Data Sources:** Ovid MEDLINE/PubMed, Embase, and PsycInfo databases were searched for studies published between January 1, 1946 and January 19, 2023 (repeated April 9, 2024, and October 15, 2024) investigating dopaminergic or serotonergic effects on reward/punishment processes in humans, according to PRISMA guidelines.

**Study Selection:** Studies reporting randomized, placebo-controlled, dopaminergic or serotonergic manipulations on a behavioral outcome from a reward/punishment processing task in healthy humans were included.

**Data Extraction and Synthesis:** Standardized mean difference (SMD) scores were calculated for the comparison between each drug (dopamine/serotonin) and placebo on a behavioral reward or punishment outcome and quantified in random-effects models for overall reward/punishment processes and four main subcategories. Study quality (Cochrane Collaboration’s tool), moderators, heterogeneity, and publication bias were also assessed.

**Main Outcome(s) and Measure(s):** Performance on reward/punishment processing tasks.

**Results:** In total, 68 dopamine and 39 serotonin studies in healthy volunteers were included (N_dopamine_=2291, N_placebo_=2284; N_serotonin_=1491, N_placebo_=1523). Dopamine was associated with an increase in overall reward (SMD=0.18, 95%CI [0.09 0.28]) but not punishment function (SMD=-0.06, 95%CI [−0.26,0.13]). Serotonin was not meaningfully associated with overall punishment (SMD=0.22, 95%CI [−0.04,0.49]) or reward (SMD=0.02, 95%CI [−0.33,0.36]). Importantly, dopaminergic and serotonergic manipulations had distinct associations with subcomponents. Dopamine was associated with reward learning/sensitivity (SMD=0.26, 95%CI [0.11,0.40]), reward discounting (SMD=-0.08, 95%CI [−0.14,-0.01]) and reward vigor (SMD=0.32, 95%CI [0.11,0.54]). By contrast, serotonin was associated with punishment learning/sensitivity (SMD=0.32, 95%CI [0.05,0.59]), reward discounting (SMD=-0.35, 95%CI [−0.67,-0.02]), and aversive Pavlovian processes (within-subject studies only; SMD=0.36, 95%CI [0.20,0.53]).

**Conclusions and Relevance:** Pharmacological manipulations of both dopamine and serotonin have measurable associations with reinforcement learning in humans. The selective associations with different components suggests that reinforcement learning tasks could form the basis of selective, mechanistically interpretable biomarkers to support treatment assignment.

**Key points:** *Question:* Do pharmacological manipulations of dopamine and serotonin affect components of reinforcement learning in humans?

*Findings:* Upregulating dopamine is associated with increased reward learning/sensitivity and reward response vigor, and decreased reward discounting. Upregulation of serotonin is associated with increased punishment learning/sensitivity and decreased reward discounting.

*Meaning:* Pharmacological manipulations of dopamine and serotonin have dissociable associations with different components of reinforcement learning. This forms a necessary basis for the development of selective markers for treatment assignment.

## Introduction

Dopamine and serotonin are two critical neuromodulators in the brain, both playing central roles in clinical and basic neuroscience. Clinically, they are the targets of different antidepressants, which are equally effective in treating mood disorders^1^. Yet, basic neuroscience suggests they serve distinct functions in behavior, particularly in reinforcement learning (RL). If dopamine and serotonin indeed have distinct effects in RL, as current theories predict, this could help identify new mechanisms to guide much-needed optimized treatments. Conversely, if they prove indistinguishable in humans, prevailing neuroscientific models would require substantial revision. Resolving this question in humans is therefore critical.

RL describes how rewards and losses guide behavior and is strongly related to depression. It is impaired in individuals who are currently depressed^2,3^, and has substantial face validity as a core mechanism leading to or maintaining depression^4^. RL is also intricately linked to the neuromodulator systems dopamine and serotonin as the brain appears to use these to broadcast key RL signals^5,6^.

However, a key challenge here lies in reconciling two seemingly contradictory bodies of research. On one hand, antidepressants targeting dopamine or serotonin appear similarly effective in treating depression^1^. On the other hand, substantial neuroscience research indicate that dopamine and serotonin play distinct roles in RL^5–8^. One possible resolution is that different antidepressants engage different components of RL. Such a selectivity could potentially explain why individuals who do not respond to serotonergic medications may respond to more dopaminergic medications. Indeed, dopaminergic antidepressants are second-line options when first-line serotonergic treatments fail^9,10^. As a necessary precondition for this, manipulating different neuromodulators must demonstrably affect distinct RL processes in humans—a question that remains unresolved. If validated, RL-based measures could serve as markers to guide treatment choices and inform the development of novel therapies. Thus, adopting an RL framework to clarify the cognitive roles of dopamine and serotonin represents a critical first step toward identifying candidate mechanisms driving the therapeutic effects of these antidepressants.

RL is however not a single process but a collection of mechanisms that are hypothesized to involve dopamine and serotonin in distinct ways^11^. Basic neuroscience highlights that dopamine is a key modulator of appetitive processes, particularly reward learning, reward discounting and reward response vigor^6,11–13^. Serotonin, while more complex, is often linked to aversive processes such as punishment learning, or inhibition in the face of punishments^8,14^. However, human studies on these neuromodulators in RL have yielded mixed and sometimes conflicting results^15,16^. These inconsistencies hinder our understanding of dopamine and serotonin in human behavior and the development of optimized treatments for mood and anxiety disorders. Therefore, we performed a meta-analysis of the effects of pharmacological manipulations of dopamine and serotonin on RL. Given the limited number of studies, for increased power we a) considered pharmacological manipulations broadly, not only antidepressants; and b) focused on healthy participants due to few patient studies. Our first aim was to ask whether dopamine and serotonin alter overall reward and punishment processing. Our second aim was to identify whether they affect specific RL subcomponents. Based on theoretical frameworks and a systematic review of the literature we identified and focused the meta-analysis on the following subcomponents of RL.

### Reward and punishment learning

A key component of RL involves trial-and-error learning from rewards and punishments via prediction errors (the difference between expected and actual outcomes)^6^. Dopamine has been extensively implicated in signaling reward prediction errors^6,17^. Some theories also postulate that dopamine dips may drive punishment learning^18^, while computational models suggest this is under serotonin’s control^14,19^. Recent animal studies have however implicated serotonin in reward learning as well, firing to reward-predicting cues or contrarily, suppressing reward learning^5,20^. To examine these processes in humans, studies mostly employ probabilistic instrumental learning tasks.

### Pavlovian bias

Choices can be driven by reflexive Pavlovian biases, where appetitive biases prompt approach toward rewards and aversive biases trigger inhibition in response to punishment, even when suboptimal^21^. Influential models propose that serotonin promotes aversive Pavlovian biases, i.e., inhibition in aversive contexts^8,22–24^. Human studies often use go/no-go tasks where these biases interfere with optimal actions or Pavlovian-instrumental transfer tasks^21,25^.

### Reward discounting

Decision-making also requires balancing rewards against costs. Current theories predict that dopamine signals benefits over costs, while serotonin may promote willingness to wait or exert effort for larger rewards by modulating perceived costs. Both mechanisms thus contribute to reduced reward discounting^26–30^. These processes are studied using tasks that contrast immediate or low effort, small rewards, with delayed or high effort, larger reward, explicit choices.

### Reward response vigor

Motivation involves deciding how vigorously actions need to be executed to obtain rewards. Dopamine is theorized to signal average reward rates, enhancing response vigor^31,32^. The role of serotonin is less defined, though an early computational theory suggest it may also modulate the long-term reward rates^14^. Tasks in this domain assess reaction times or effort exertion relative to reward magnitude or average reward rate.

Overall, this meta-analysis aims to provide a clearer understanding of how dopamine and serotonin shape RL in humans, contributing to both theoretical and clinical advances.

## Methods

### Systematic review

We conducted a preregistered systematic review and meta-analysis (PROSPERO:CRD42022363747) according to PRISMA guidelines^33^ (eAppendix in the Supplement). The Ovid MEDLINE/PubMed, Embase, and PsycInfo databases were searched for articles published between January 1, 1946, and January 21, 2023. The search was repeated on April 9, 2024 and October 15, 2024 for additional articles published since 2023. Titles or abstracts were searched containing the terms (dopamine* or seroton* or tryptophan* or tyrosine*) and (reward* or motivat* or punish* or reinforce* or decision* or effort* or learn* or incentive* or volition or choice*) and (participa* or subject* or individ* or human* or healthy* or investigat* or experiment*) and restricted to articles in English and humans. Detailed description of the search strategy is described in eTable1.

We included studies reporting 1) interventions manipulating either dopamine or serotonin, in 2) healthy human volunteers (18-65 years old), 3) randomized placebo-controlled studies that 4) reported a behavioral outcome from a reward and/or punishment processing task. We only included samples that 5) did not overlap with other included datasets and 6) contained sufficient information to calculate standardized mean difference (SMD) scores between placebo and drug conditions (see Supplement).

### Meta-analysis

Effect sizes were calculated for behavioral outcomes relating to reward and punishment learning/sensitivity, aversive and appetitive Pavlovian bias, reward vigor, and reward discounting. To evaluate overall reward and punishment processes, we consolidated the components into two main categories. The overall reward category combined reward learning/sensitivity, appetitive Pavlovian bias, reward vigor, and reward discounting–reversing the discounting direction so that a positive SMD represents increased reward weighting. The overall punishment function combined punishment learning/sensitivity and aversive Pavlovian bias. We also identified risk attitude and model-based learning/flexibility as two additional RL subcomponents. However since these are not straightforward to combine with the other subcomponents, we consider them separately (see Supplement).

Pharmacologically, we pooled data from both agonists and antagonists, assuming symmetrical effects for drugs that upregulate or downregulate a neuromodulator. We recoded antagonist effects as if they were agonist effects. For example, if a study reported that a dopamine antagonist reduced reward vigor, we recoded this effect to represent an agonist increasing reward vigor. Some dopamine studies used single low doses of agonists/antagonists intended to specifically target presynaptic autoreceptors. These effects were interpreted contrary to their typical activity profile (e.g., a low-dose agonist was coded as an antagonist), as suggested by the literature (see Supplement). Thus, for a low-dose dopamine agonist that e.g., reduced reward learning, we first classified it as an antagonist reducing reward learning, and then recoded it as an agonist increasing reward learning, maintaining a consistent directionality across studies. To assess the impact of this interpretation, we conducted sensitivity analyses using the original drug activity profiles as well.

Both within- and between-subject designs were included. The detailed procedures for calculating between- and within-subject SMDs including the conversion procedures are described in Supplement, with comparisons of different effect size conversions for within-subject studies depicted in eFigure1. Random-effects models were conducted in R (version 4.3.3) using metafor^34^ packages to examine the association between upregulating dopamine and serotonin with placebo on overall reward and punishment processes and subcomponents. Heterogeneity, publication bias, moderators and study quality were also assessed (see Supplement).

## Results

Data from 107 studies were included (N_dopamine_=2291, N_placebo_=2284; N_serotonin_=1491, N_placebo_=1523; Figure 1; eTable2). Risk of bias was generally high for both randomization and allocation concealment, but low for blinding of outcome assessment (eFigure2).

**Figure 1:**
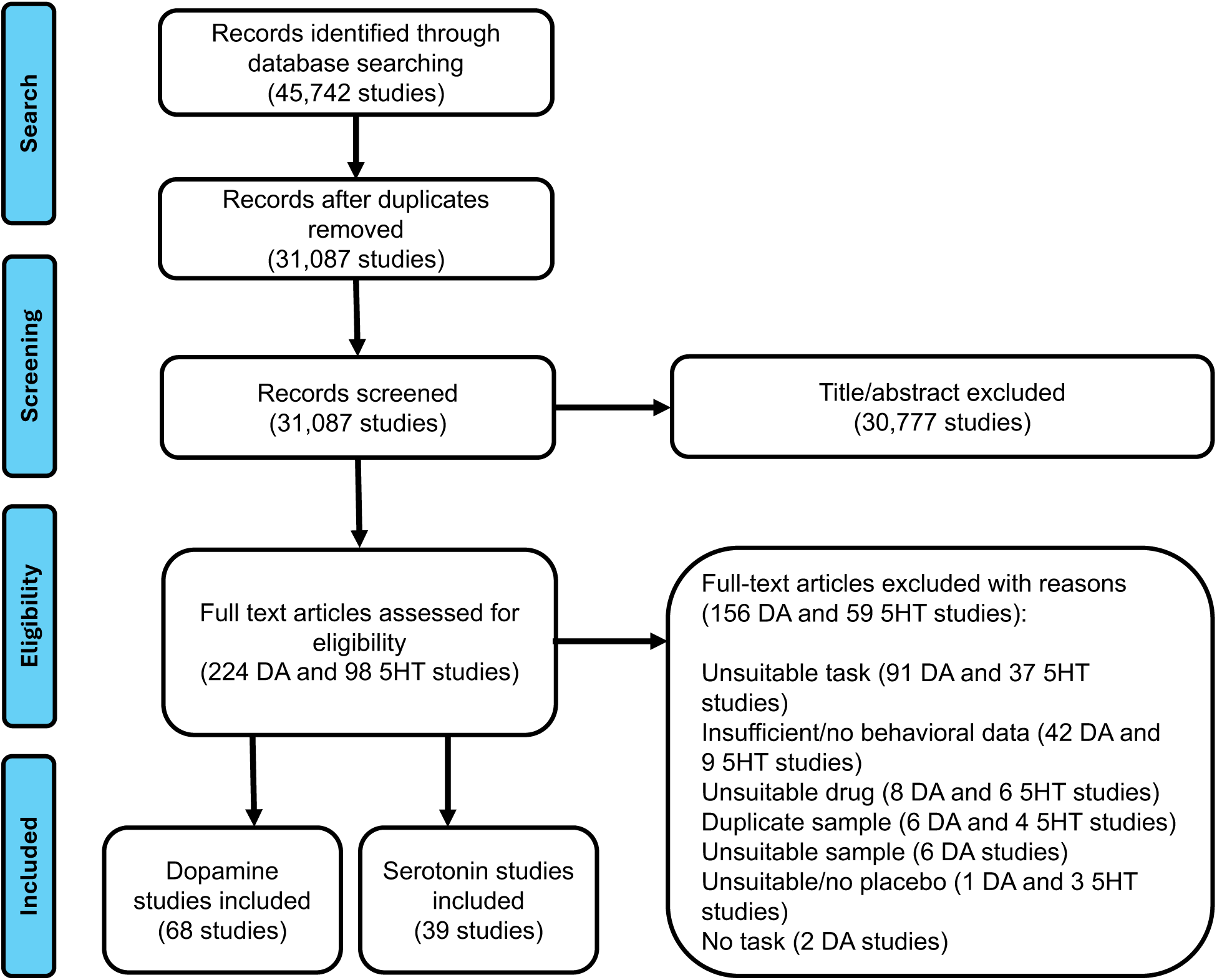
Flow Diagram of Study Selection and Inclusion. Abbreviations: DA: dopamine; 5HT: serotonin.

### Meta-analysis results

#### Dopamine

Overall, dopamine had a small positive association with reward processes (SMD=0.18, 95%CI [0.09, 0.28]) but a negligible negative association with overall punishment processes (SMD=-0.06, 95%CI [−0.26, 0.13]; eFigure3). Analyses according to the original drug activity profile changed inference for the overall reward process; eFigure4).

Dopamine increased reward learning/sensitivity (Figure 2a; SMD=0.26, 95%CI [0.11, 0.40]), reward vigor (SMD=0.32, 95%CI [0.11, 0.54]; Figure 2c) and decreased reward discounting (SMD=-0.08, 95%CI [−0.14, −0.01]; Figure 2b), but were not associated with any other subcomponents (Figure 3; eFigure5). Analyses according to the original drug activity profile influenced the inference of dopamine on reward learning/sensitivity and reward discounting but no other ones (eFigure6). We also explored the associations with computational learning rate and sensitivity parameters, but there were far fewer studies reporting these and no clear pattern emerged (eFigure7).

**Figure 2:**
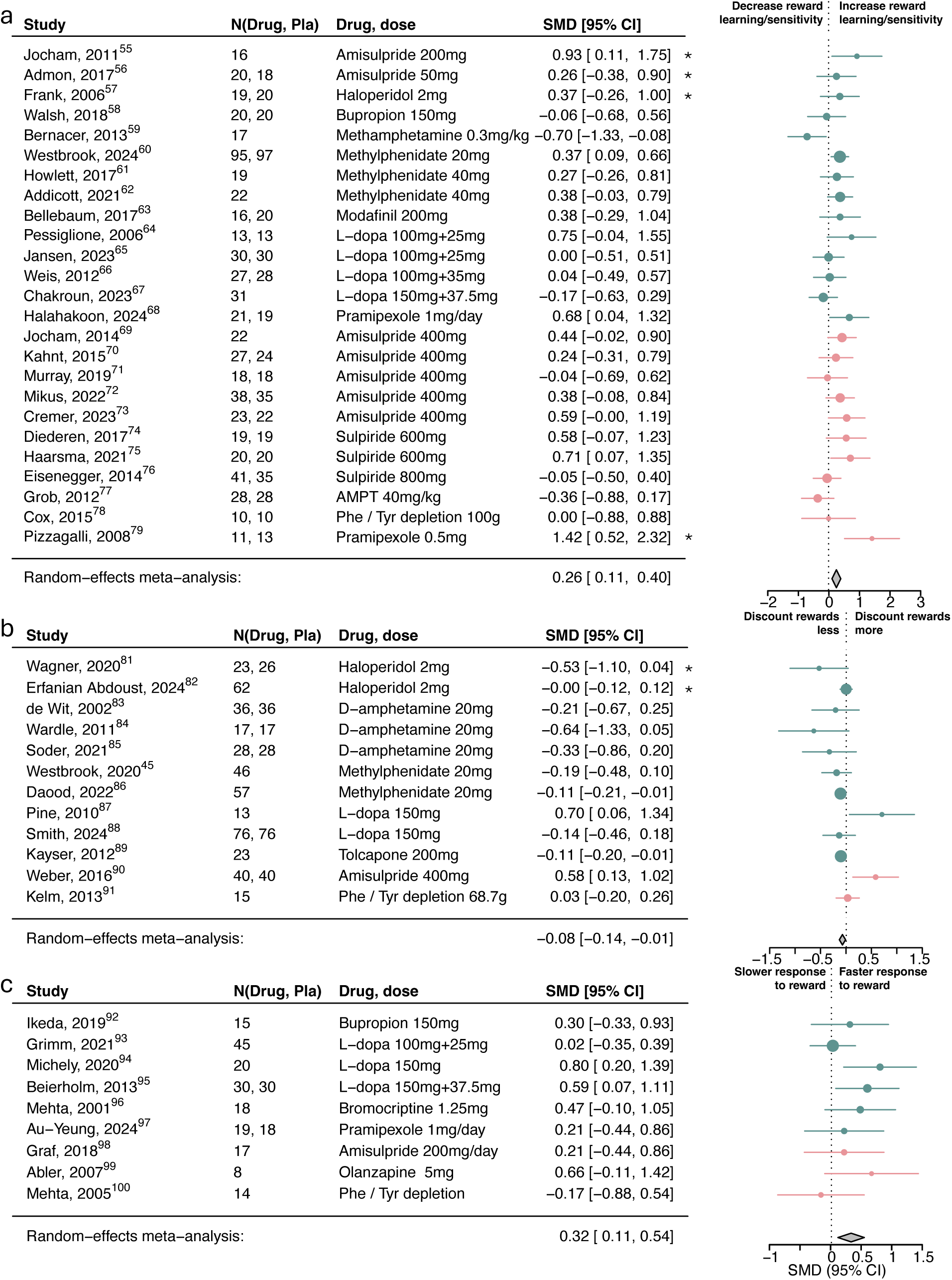
Association of dopamine upregulation with reward subcomponents. Standardized mean differences (SMDs) of the association between upregulating dopamine versus placebo and a) reward learning/sensitivity, b) reward discounting, and c) reward response vigor. Pla indicates placebo. We recoded antagonist effects as if they were agonists (pink SMD point estimates and 95% CI). Green SMD point estimates and 95% CI indicate studies that were coded as original agonists. Asterisks indicate studies that used a low dose of an agonist or antagonists. These effects were interpreted contrary to their typical activity profile (e.g., a low-dose agonist acting antagonistically), as suggested by the literature.

**Figure 3:**
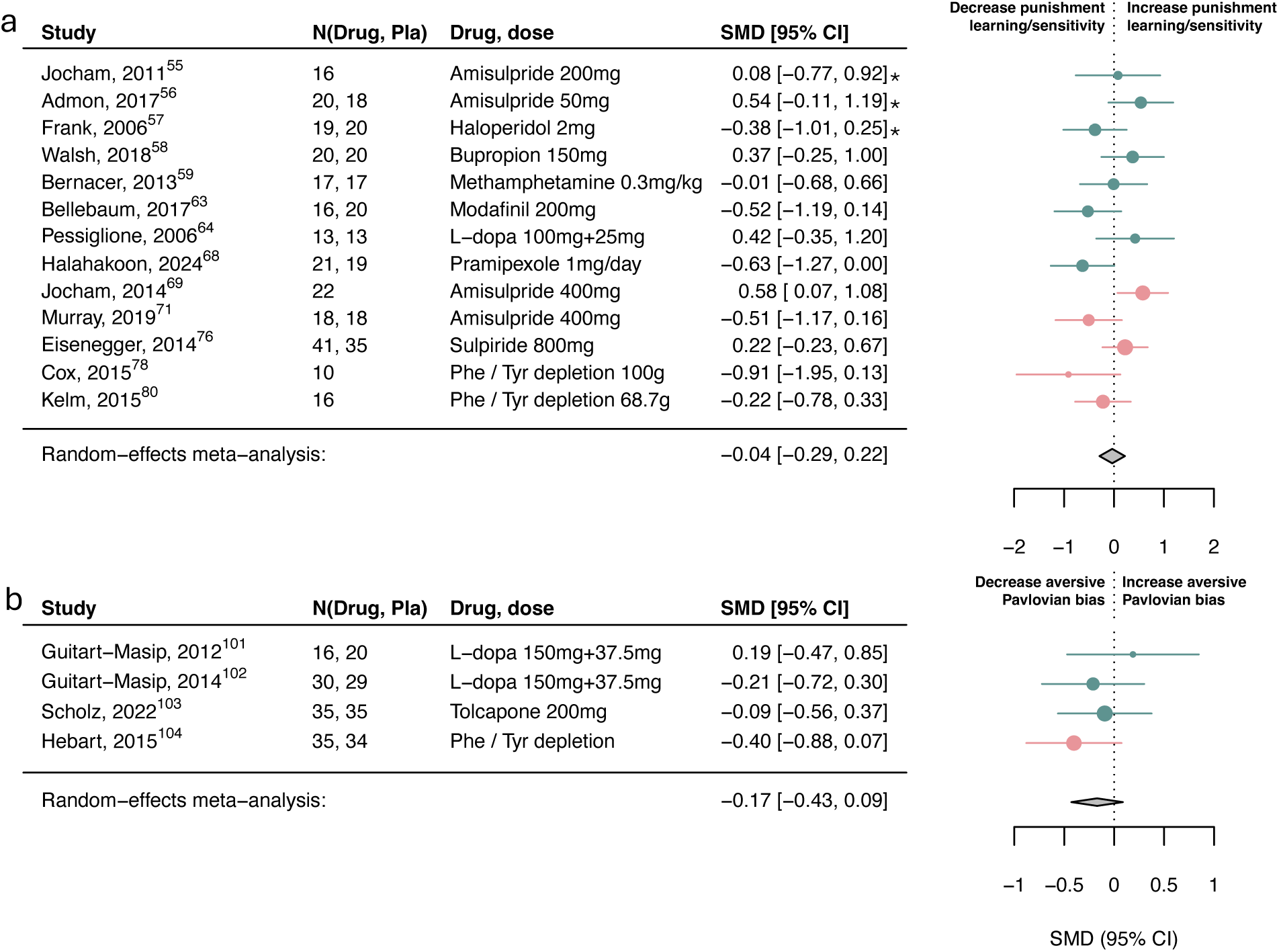
Association of dopamine upregulation with punishment subcomponents. Standardized mean differences (SMDs) of the association between upregulating dopamine versus placebo and a) punishment learning/sensitivity and b) aversive Pavlovian bias. We recoded antagonist effects as if they were agonists (pink SMD point estimates and 95% CI). Green SMD point estimates and 95% CI indicate studies that were coded as original agonists. Asterisks indicate studies that used a low dose of an agonist or antagonists. These effects were interpreted contrary to their typical activity profile (e.g., a low-dose agonist acting antagonistically), as suggested by the literature.

There was low-to-moderate heterogeneity across all overall processes and main subcomponents (see Supplement). Substantial heterogeneity was only found in the risk attitude and model-based/flexibility subcomponents, with outlier exclusion resolving risk attitude heterogeneity, resulting in a significant positive dopamine association (eFigure8).

#### Serotonin

Overall, serotonin did not show associations with reward (SMD=0.02, 95%CI [−0.33, 0.36]) or punishment processes (SMD=0.22, 95%CI [−0.04, 0.49]; eFigure9).

Serotonin had a small-to-moderate association with reward discounting (SMD=-0.35, 95%CI [−0.67, −0.02]; Figure 4c) and punishment learning/sensitivity (SMD=0.32, 95%CI [0.05, 0.59]; Figure 5a) but not with reward learning/sensitivity, reward vigor, Pavlovian biases (Figure 4-5) or other subcomponents (eFigure10).

**Figure 4:**
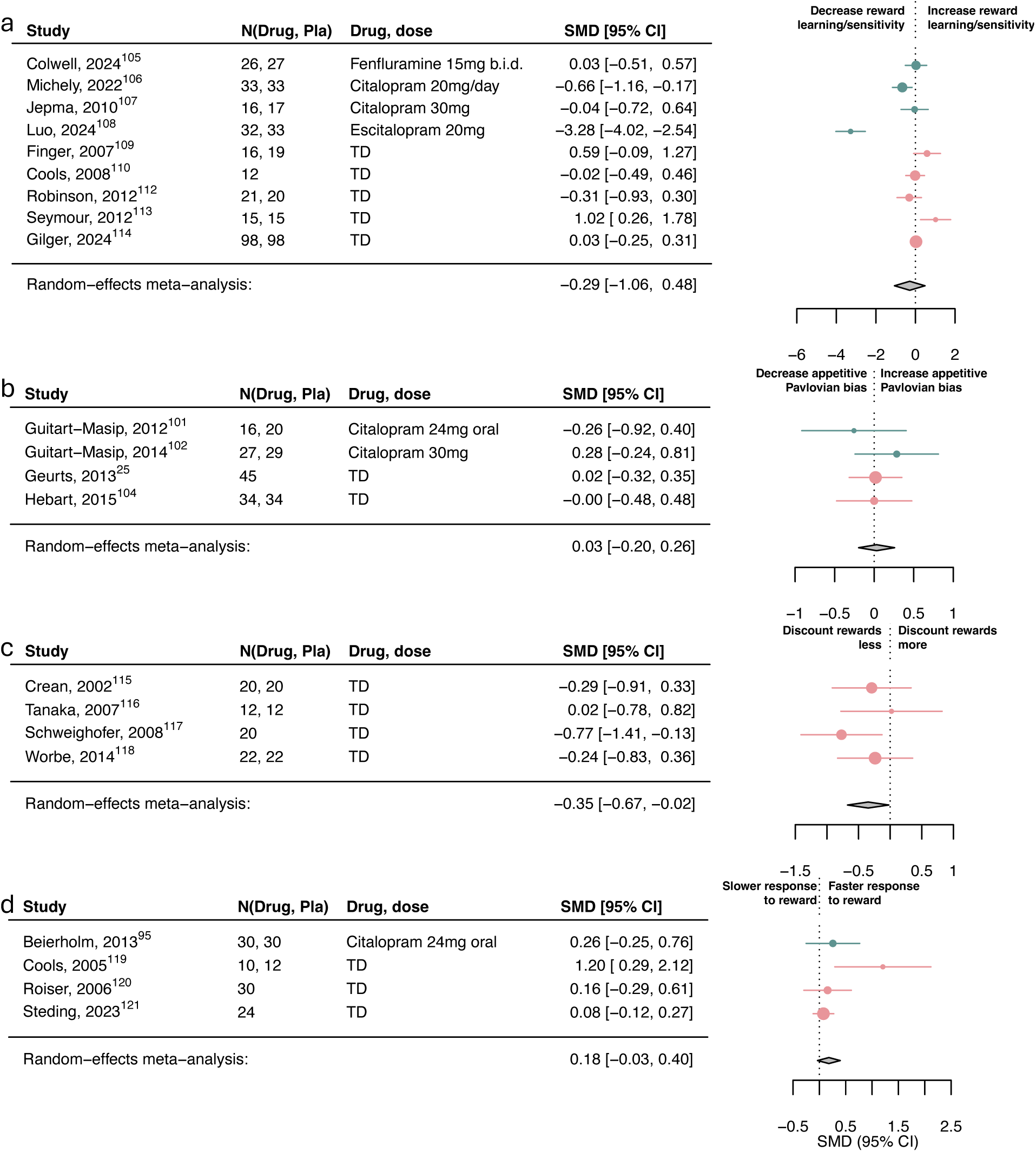
Association of serotonin upregulation with reward subcomponents. Standardized mean differences (SMDs) of the association between upregulating serotonin versus placebo and a) reward learning/sensitivity, b) appetitive Pavlovian bias, c) reward discounting, and d) reward response vigor. Pla indicates placebo. We recoded antagonist effects as if they were agonists (pink SMD point estimates and 95% CI). Green SMD point estimates and 95% CI indicate studies that were coded as original agonists. TD: tryptophan depletion.

**Figure 5:**
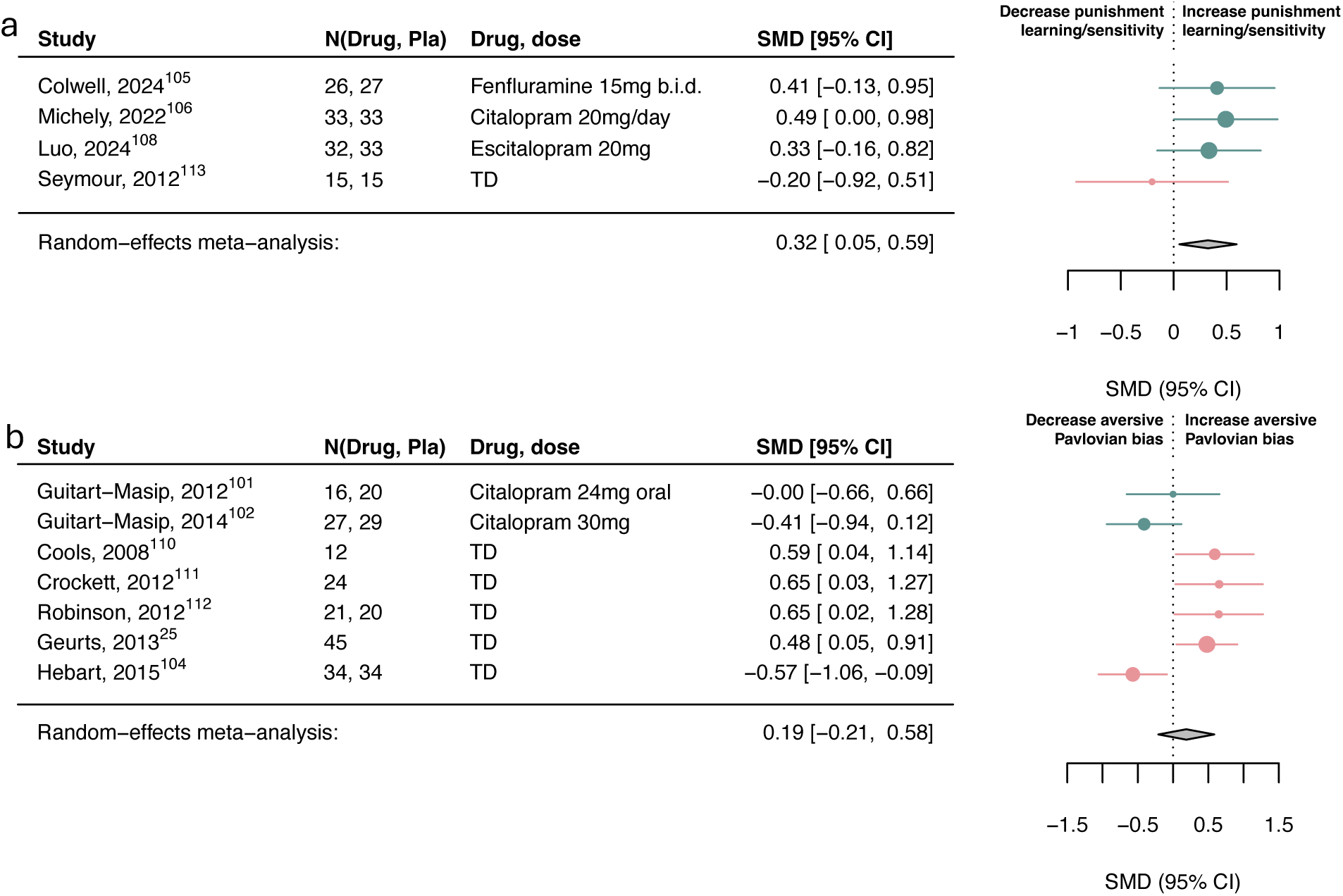
Association of serotonin upregulation with punishment subcomponents. Standardized mean differences (SMDs) of the association between upregulating serotonin versus placebo and a) punishment learning/sensitivity and b) aversive Pavlovian bias. We recoded antagonist effects as if they were agonists (pink SMD point estimates and 95% CI). Green SMD point estimates and 95% CI indicate studies that were coded as original agonists. TD: tryptophan depletion.

However, three within-subject studies had to be excluded in the aversive Pavlovian domain due to missing information needed to combine with between-subject studies. Examining the six within-subject studies separately, serotonin was associated with a small-to-medium increase in aversive Pavlovian processes (SMD=0.36, 95%CI [0.20, 0.53]; eFigure11). Exploring learning rate and sensitivity computational parameters, serotonin was associated with increased punishment learning rates and decreased punishment sensitivity (eFigure12). However, this finding is based on a limited number of studies.

There was substantial heterogeneity in overall reward/punishment categories and the reward learning/sensitivity and aversive Pavlovian subcomponents. Removing identified outliers explained some of this variance and revealed a significant increase of overall punishment processes by serotonin (SMD=0.32, 95%CI [0.10, 0.54]; see Supplement; eFigure13).

### Publication bias and moderator analyses

Egger’s test suggested significant publication bias in the overall dopamine reward domain (z=2.28, p=0.02) but not in any of the other overall domains (dopamine punishment: z=-0.79, p=0.43; serotonin reward: z=0.07, p=0.94; serotonin punishment: z=-0.09, p=0.93; eFigure14).

#### Dopamine

Only the reward and punishment learning/sensitivity and reward discounting categories contained greater than 9 studies, allowing a test for funnel plot asymmetry^35^. None of these showed significant publication bias (reward learning/sensitivity: z=1.49, p=0.14; punishment learning/sensitivity: z=-1.42, p=0.15; reward discounting: z=0.057, p=0.95; eFigure15).

Drug type significantly moderated the reward discounting results (R^2^=13.88%, I^2^=6.45%, p=0.03), with original agonist studies decreasing reward discounting (SMD=-0.10, 95%CI [−0.16, - 0.03]; 10 studies), while studies that originally used antagonists showed no meaningful association (SMD=0.14, 95%CI [−0.06, 0.35]; 2 studies). None of the other moderators had an effect (see Supplement).

#### Serotonin

The model-based learning/flexibility subcomponent was the only one with sufficient studies to conduct a funnel plot asymmetry analysis, which revealed no significant asymmetry (z=1.45, p=0.15; eFigure15).

No meaningful moderators were identified for serotonin (see Supplement).

## Discussion

This meta-analysis reveals distinct roles for dopamine and serotonin in modulating RL processes in humans. Dopamine increased reward learning and sensitivity, and reward vigor. A different pattern of effects was observed for serotonin: promoting learning from punishments, and possibly amplifying aversive Pavlovian biases. Both serotonin and dopamine increased the value of rewards when contrasted with costs of time or effort. These findings contribute to two key domains: advancing neuroscience theories of dopamine and serotonin in humans, and informing clinical applications. We begin by addressing the theoretical implications and conclude with the clinical relevance.

The differing associations of dopamine and serotonin are broadly consistent with theoretical expectations. The most influential theory of dopamine posits that dopamine signals reward prediction errors, driving reward learning^6,36^. Supporting this, we found that boosting dopamine promotes reward learning/sensitivity in humans. Unlike preclinical studies, however^5,20^, we found no serotonin association with reward learning, reinforcing the idea that dopamine and serotonin serve dissociable functions here. Instead, serotonin was selectively associated with punishment learning, consistent with theories and human research linking serotonin to aversive processing^8,37^, and highlighting an opposing role to dopamine in learning^14^.

We also provide preliminary evidence for a dissociation of serotonin and dopamine in Pavlovian biases. Specifically, serotonin—but not dopamine—was associated with enhanced aversive biases in within-subject studies. This resonates with findings in depressed patients treated with serotonergic antidepressants^38^, and supports theories suggesting serotonin facilitates inhibition specifically in punishment contexts^8,22–24^. However, this effect is not consistently observed, potentially owing to high measurement noise in these tasks^39,40^, underscoring the need for psychometrically optimized tasks^41^.

Beyond learning, it is noteworthy that we found dopamine influencing reward vigor and discounting. This aligns with animal work suggesting that dopamine encodes a unified motivational signal, influencing both how energetically actions are taken (vigor) and decisions of whether to invest in effortful activity (value-based choice)^32,42^. At present, it is unclear whether the dopamine associations with vigor and discounting (especially effort-based discounting which supplementary exploratory analyses indicated might be particularly associated with dopamine) are related in humans as well. Notably, however, our findings converge with a rodent and clinical meta-analysis, where dopaminergic agents affected both reward discounting and vigor^32,42–44^.

Like dopamine, serotonin also diminished reward discounting. Although this might suggest that both have similar functions here, a more likely explanation is that they affect reward discounting through distinct mechanisms. Prior research suggests that dopamine reduces discounting by boosting the perceived benefits while serotonin may alter the perceived costs^27,28,45^. Although we were unable to dissociate the precise mechanism here, it is interesting that the serotonin studies exclusively covered delay discounting tasks (while dopamine additionally covered effort discounting tasks). This may suggest that serotonin increases the discounting factor in RL models to promote a focus on long-term over short-term outcomes^27,46^.

Clinically, perhaps the most significant finding is that dopaminergic and serotonergic pharmacological manipulations had appreciably distinct associations. They engaged different components of RL in a manner which is measurable and identifiable with simple behavioral tasks. Hence, simple RL tasks may play a useful role in measuring treatment target engagement. This contrasts with symptom markers, which do not differentiate psychiatric medications^47,48^. Interestingly, our observed effect sizes, though modest, align with RL impairments observed in mood disorders^2,3^, dopamine effects in Parkinson’s^44^, and antidepressant efficacy^1^. Our results thus fulfill a key precondition for future work: if different antidepressants modulate distinct RL components, then RL-based biomarkers could guide more optimized treatments. This aligns with calls to use RL to understand antidepressant mechanisms and their clinical relevance^49^. While individual-level precision requires further refinement, establishing, even subtle, systematic group-level neuromodulatory RL associations is a crucial first step in guiding those efforts. A key and important question for future work is whether individual differences in baseline RL measures could help identify individuals more likely to benefit from one or the other type of intervention. This will require studies which simultaneously measure multiple RL domains, psychometrically robust tasks and direct head-to-head comparison of the treatments of interest.

### Limitations

Several limitations warrant consideration. First, few studies reported computational RL outcomes, forcing us to rely on broad measures. While we attempted to examine learning rate and sensitivity separately, this proved challenging as few reported these. It is further hard to disentangle them (indeed, serotonin increased punishment learning but decreased punishment sensitivity; eFigure12) and both are sensitive to task designs^50^. Despite this, meaningful variability is evident in broad measures.

Second, the lack of studies directly comparing dopamine and serotonin prevented us from examining this relationship. However, such comparisons will be crucial for drawing definitive conclusions about their differential effects. There is also a lack of pharmacological specificity in the dopamine domain. Some medications target both dopamine and noradrenaline, complicating interpretation.

Third, the association between dopamine and reward learning and discounting depended on the drug-binding interpretation, shifting from negligible to meaningful when low-dose presynaptic accounts were considered. This highlights the complexities of dopaminergic research, including evidence that the direction of effects may depend on baseline dopamine levels^51^. Such complexities might therefore have added noise to the effect size estimates.

Fourth, substantial heterogeneity was observed in the association between serotonin and reward learning, potentially due to the lack of common tasks across studies. This suggests the meta-analytic effect may be fragile, underscoring the need for improved task standardization and psychometrically robust tasks. Several other factors may have contributed to heterogeneity, such as feedback saliency^52^ or menstrual cycle^53^, which future research needs to assess.

Fifth, most serotonin studies used tryptophan depletion (TD), a method debated as a mild serotonin manipulation^54^. However, our findings show meaningful associations even in TD-only analyses (reward discounting, aversive Pavlovian processes), supporting their validity despite this debate.

We also pooled within- and between-subject data to increase power, applying effect size corrections of within-subject studies where feasible. When not possible, inflated variance of effect sizes would be the likely consequence, reducing our power to detect significant effects.

Finally, all studies were done in healthy controls. Ultimately, however, head-to-head comparisons in patients across tasks will be required to establish the utility of RL processes for clinical applications.

## Conclusions

Despite methodological heterogeneity, there is evidence that dopamine and serotonin affect RL; and that they do so by altering specific, distinct processes. These insights are valuable as they set forth candidate mechanisms for understanding the differentiating effects of treatments targeting different neuromodulators. Critically, the selective associations are observable in highly scalable human behavioral tasks, suggesting that such assessments may provide a fertile ground for the development of differentially sensitive biomarkers.

## Acknowledgments

We thank the Applied Computational Psychiatry group for helpful comments and discussion on an earlier draft of the manuscript. We also thank the authors who provided raw data for the articles in these analyses. Some of these data were provided (in part) by the Radboud University, Nijmegen, The Netherlands. For the purpose of Open Access, the author has applied a CC BY public copyright license to any Author Accepted Manuscript version arising from this submission. This research was funded by Wellcome Trust grants (221826/Z/20/Z and 226790/Z/22/Z) and a Carigest S.A. donation to QJMH. We obtained support by the UCLH NIHR BRC. MB was supported by the Office for Life Sciences and the National Institute for Health and Care Research (NIHR) Mental Health Translational Research Collaboration, hosted by the NIHR Oxford Health Biomedical Research Centre and by the NIHR Oxford Health Biomedical Research Centre. The views expressed are those of the authors and not necessarily those of the UK National Health Service, the NIHR, or the UK Department of Health and Social Care. The funders had no role in the design and conduct of the study; collection, management, analysis, and interpretation of the data; preparation, review, or approval of the manuscript; and decision to submit the manuscript for publication. AM and QJMH had full access to all the data in the study and take responsibility for the integrity of the data and the accuracy of the data analysis.

## Additional information

The preregistration can be found at https://www.crd.york.ac.uk/PROSPERO/display_record.php?RecordID=363747 and open data and code to replicate all analyses are available online: https://github.com/huyslab/DA5HT_MetaAnalysis_public.

