## Supplementary Materials for "Dopamine and serotonin differentially associated with reward and punishment processes in humans: A systematic review and meta-analysis"

#### Table of Contents

*1 Supplementary Methods* ... 2

1.1 Search strategy ... 2

1.2 Paper selection ... 4

1.3 Meta-analysis ... 5

1.3.1 Risk attitude and model-based learning/flexibility ... 5

1.3.2 Categorization of dopaminergic agents ... 6

1.3.3 Assessment of heterogeneity and outliers ... 7

1.3.4 Assessment of publication bias and moderators ... 7

1.3.5 Assessment of study quality ... 7

1.4 Effect size calculations ... 8

1.4.1 Conversion of outcome measures into effect sizes and associated variances ... 8

*2 Supplementary results* ... 13

2.1 Assessment of study quality ... 14

2.2 Additional dopamine results ... 15

2.3 Dopamine heterogeneity ... 20

2.4 Additional serotonin results ... 22

2.5 Serotonin heterogeneity ... 25

2.6 Moderator analyses ... 27

2.6.1 Dopamine ... 27

2.6.2 Serotonin ... 28

2.7 Publication bias ... 30

2.8 Additional supplementary analyses ... 32

*3 References* ... 39

*4 eAppendix 1* ... 50

4.1 PRISMA 2020 Checklist ... 50

4.2 PRISMA 2020 Abstract Checklist ... 53

### 1 Supplementary Methods

#### 1.1 Search strategy

eTable 1 lists our search strategy for each database via Ovid, including the MeSH terms. The search terms were refined on a set of 40 in-lab references[1-40] to ensure they captured all relevant studies.

| | MEDLINE/PubMed | | Embase | | PsycInfo |
|---|---|---|---|---|---|
| 1 | Dopamine/ | 1 | catecholamine/ | 1 | exp catecholamines/ |
| 2 | exp Dopamine Agents/ | 2 | exp *dopamine receptor stimulating agent/ | 2 | exp dopamine agonists/ |
| 3 | Serotonin/ | 3 | exp *serotonin receptor affecting agent/ | 3 | exp dopamine antagonists/ |
| 4 | exp Serotonin Agents/ | 4 | tryptophan/ | 4 | exp Serotonin Agonists/ or exp Serotonin/ or exp Serotonin Antagonists/ |
| 5 | tryptophan/ or tyrosine/ | 5 | tyrosine/ | 5 | tryptophan/ |
| 6 | dopamine*.ti,ab. | 6 | dopamine*.ti,ab. | 6 | tyrosine/ |
| 7 | seroton*.ti,ab. | 7 | seroton*.ti,ab. | 7 | dopamine*.ti,ab. |
| 8 | tryptophan*.ti,ab. | 8 | tryptophan*.ti,ab. | 8 | seroton*.ti,ab. |
| 9 | tyrosine*.ti,ab. | 9 | tyrosine*.ti,ab. | 9 | tryptophan*.ti,ab. |
| 10 | 1 or 2 or 3 or 4 or 5 or 6 or 7 or 8 or 9 | 10 | 1 or 2 or 3 or 4 or 5 or 6 or 7 or 8 or 9 | 10 | tyrosine*.ti,ab. |
| 11 | learning/ or avoidance learning/ or reinforcement, psychology/ or punishment/ or reward/ or reversal learning/ | 11 | learning/ or exp *associative learning/ or exp *conditioning/ or exp *"reinforcement (psychology)"/ or exp *reversal learning/ or exp *"reinforcement learning (machine learning)"/ | 11 | 1 or 2 or 3 or 4 or 5 or 6 or 7 or 8 or 9 or 10 |
| 12 | motivation/ or exploratory behavior/ or goals/ | 12 | exp *monetary reward/ or reward seeking behavior/ or reward/ | 12 | learning/ or reward learning/ |
| 13 | decision making/ or choice behavior/ or delay discounting/ or uncertainty/ | 13 | exp *motivation/ or exploratory behavior/ | 13 | behavior/ or exp approach avoidance/ or exp approach behavior/ or exp avoidance/ or exploratory behavior/ |
| 14 | (reward* or motivat* or punish* or reinforce* or decision* or effort* or learn* or incentive* or choice*).ti,ab. | 14 | exp *decision making/ or exp *cognitive bias/ | 14 | computational reinforcement learning/ |
| 15 | 11 or 12 or 13 or 14 | 15 | (reward* or motivat* or punish* or reinforce* or decision* or effort* or learn* or incentive* or choice*).ti,ab. | 15 | exp reinforcement/ |
| 16 | (participa* or subject* or individ* or human* or healthy* or investigat* or experiment*).ti,ab. | 16 | 11 or 12 or 13 or 14 or 15 | 16 | exp motivation/ |
| 17 | research subjects/ or healthy volunteers/ or volunteers/ | 17 | (participa* or subject* or individ* or human* or healthy* or investigat* or experiment*).ti,ab. | 17 | exp decision making/ |

| | | | | | |
|---|---|---|---|---|---|
| 18 | 16 or 17 | 18 | research subject/ or volunteer/ or normal human/ | 18 | exp decision theory/ |
| 19 | 10 and 15 and 18 | 19 | 17 or 18 | 19 | exp delay discounting/ |
| 20 | limit 19 to (english language and "humans only (removes records about animals)") | 20 | 10 and 16 and 19 | 20 | (reward* or motivat* or punish* or reinforce* or decision* or effort* or learn* or incentive* or choice*).ti,ab. |
| | | | | 21 | 12 or 13 or 14 or 15 or 16 or 17 or 18 or 19 or 20 |
| | | | | 22 | (participa* or subject* or individ* or human* or healthy* or investigat* or experiment*).ti,ab. |
| | | | | 23 | exp experimental subjects/ |
| | | | | 24 | volunteers/ |
| | | | | 25 | 22 or 23 or 24 |
| | | | | 26 | 11 and 21 and 25 |
| | | | | 27 | limit 26 to (human and english language) |

**eTable 1: Complete search and MeSH terms for each database.**

#### 1.2 Paper selection

Data were requested from authors mainly to obtain sufficient information from within-subject studies to pool together with between-subject studies or to obtain individual data for high and no/low reward conditions in the reward response vigor category. If a study reported data from multiple tasks, we extracted data from the most commonly used task in that category. We also prioritized parameters over model-agnostic measures wherever possible, and outcome measures that were as similar to other outcome measures within the same category.

In cases where multiple drugs or dosages were reported, we chose the more commonly used drug in that category and the highest dosage, respectively. For example, for studies that included both tryptophan depletion and tryptophan loading conditions (N=3), we prioritized the depletion condition, as it was the more commonly used and established manipulation in the literature. This decision also ensured statistical independence, as including both conditions from the same study would introduce duplicate samples.

We excluded studies using drugs that have unclear, complex or substantially mixed dopamine or serotonin targets and/or those that would not be considered typical neuromodulatory targets of antidepressants, including LSD studies. While LSD has shown to have interesting effects on RL[41-43], LSD is considered a novel, experimental antidepressant, which is thought to act differently to traditional antidepressants and shown to act on both serotonin and dopamine receptors[44]. A key aim of this meta-analysis was to 1) focus on research that more directly relates to typical serotonergic and dopaminergic-based antidepressants and 2) examine serotonin and dopamine systems separately. Given the already high heterogeneity in our included studies, we opted to exclude psychedelics like LSD to maintain a clearer mechanistic focus on traditional serotonergic and dopaminergic modulation.

For the reward learning/sensitivity category, we only included measures that related to reward processing and could be dissociated from punishment if those were used and vice versa for the punishment learning/sensitivity category. As our initial aim was to examine RL components, we extracted computational parameters for our outcome measure where possible. For the learning/sensitivity subcomponents, we prioritized learning rate over sensitivity parameters. Sensitivity parameters are harder to interpret, as they capture both sensitivity and nonspecific noisiness. For this reason — and due to the greater number of studies using learning rates — we prioritized learning rates in our main analysis.

For the reward response vigor category, we primarily included measures that compared high-reward conditions to no-reward or low-reward conditions. This approach was intended to specifically capture the influence of dopamine/serotonin on reward-induced invigoration, rather than general motor functions. If these measures were not extractable however, we included a measure of vigor on high reward only if the effect of drug on the no or low reward condition was nonsignificant (N=2 in the dopamine domain). Excluding these two studies[45,46] slightly reduced the pooled effect size in the reward response vigor subcategory (SMD=0.23; 95% CI [0.01, 0.44]) and the overall reward domain meta-analysis for dopamine (SMD=0.16; 95% CI [0.07, 0.25]) but did not change inference.

For the overall reward/punishment meta-analyses, we excluded any studies with duplicate samples. This issue only arose in the dopamine domain of the overall reward meta-analysis, where two pairs of studies had overlapping samples[38,47,48]. As a conservative approach, we retained the studies with the smallest effect sizes from each pair.

Due to our wide search strategy, a number of studies were excluded based on unsuitable cognitive tasks. Overall, we excluded any tasks that could not be easily categorized within the RL subcomponents outlined in the introduction. Specifically, we excluded: tasks where reward and punishment learning could not be disentangled in the outcome measure (e.g., Iowa Gambling Task); tasks that did not primarily assess a reward- or punishment-based process, such as cognitive control, impulsivity, memory, perceptual, or emotional processing tasks; tasks designed primarily to study social behavior (e.g., Dictator Game, Trust Game) or drug-seeking behavior (e.g., drug-preference tasks); studies that used inappropriate outcome measures, such as those that did not report reward- or punishment-specific learning outcomes or lacked behavioral outcome measures (e.g., some Pavlovian tasks that measured only physiological responses); and tasks that were not comparable to others included in the meta-analysis, or where rewards and punishments were not explicitly used as outcome measures.

Preprints were included but abstracts or unpublished work were not. Articles were independently assessed by A.M. and either Z.Q. or T.E. Conflicts were resolved through in-person discussion with Q.J.M.H. Data were extracted by A.M. and Z.Q. and double-checked by A.M. and Z.Q. or Q.D. or Y.A.

#### 1.3 Meta-analysis

##### 1.3.1 Risk attitude and model-based learning/flexibility

As supplementary analyses, we examined risk attitude and model-based learning/flexibility. The risk attitude component examines how risk averse or risk-seeking individuals are. This component is usually examined with gamble tasks where individuals make a choice between a gamble of some risk between winning or losing and a sure or lower risk option with lower rewards.

Model-based learning/flexibility were composed of tasks that traditionally examine model-based learning or reversal learning. Model-based learning describes how individuals need to acquire a model of the task to make goal-directed choices, in contrast to model-free learning which relies on trial-and-error learning[49]. Similarly, in reversal learning tasks individuals are exposed to changes in the task rules and have to infer when the context has changed to make appropriate decisions. These tasks are thought to measure flexibility, but also model-based aspects since an understanding of the task rules are required.

Risk attitude was excluded from the main meta-analyses because, while it is categorized as a cost-benefit decision-making process and sometimes grouped with effort and temporal delay discounting, it does not necessarily integrate well with these constructs. For instance, in risk attitude paradigms, impulsivity manifests as greater risk-seeking behavior, which however corresponds to lower reward discounting by risk. In contrast, temporal delay tasks define impulsivity as choosing smaller, sooner rewards over larger, delayed ones, which also reflects higher reward discounting by time. Given these opposing directions in reward discounting when considering impulsivity and the frequent inclusion of both rewards and punishments in gambles in risk attitude tasks, we treated risk attitude as a separate construct.

The model-based learning/flexibility subcomponent was also considered as a separate construct as it contains measures that tap into a more complex reward processing domain and some outcome measures in this task had mixed reward and punishment features. We also examined within-subject studies only of both of these subcomponents since a number of within-subject studies could not be combined with the between-subject studies.

##### 1.3.2 Categorization of dopaminergic agents

For the primary analysis, we categorized low-dose dopamine agonists and antagonists based on their opposing pharmacological effects. This was based on several lines of evidence suggesting that pre- or post-synaptic binding depends on dosage. For example, low dose of D2-receptor antagonists has shown to block presynaptic autoreceptors, which are inhibitory, and therefore increase, instead of block, dopamine transmission[50-52]. Similarly, studies in humans have shown that a low dose antagonist increase ventral striatal response to rewards – a typical response to boosting dopamine[53] – as well as serum prolactin levels[14]. A similar pattern has been observed for some agonists at low dosages, like pramipexole[27].

##### 1.3.3 Assessment of heterogeneity and outliers

Heterogeneity was assessed with $\tau^2$, quantifying the variance of the true effect sizes, and the $I^2$ statistic, representing the percentage of variability in the effect sizes not accounted for by sampling error[54]. Heterogeneity was interpreted as low (25-50%), medium (50-75%) or high (>75% $I^2$)[54]. Studies were classified as outliers if their 95% confidence interval did not overlap with the pooled effect CI, using the dmetar package in R[55]. To evaluate the influence of outliers, sensitivity analyses were conducted by recalculating the pooled effect and heterogeneity after removing the outliers.

##### 1.3.4 Assessment of publication bias and moderators

Small-study effects were assessed with the Egger test[56] ($p<0.05$) for categories with at least 10 studies[57]. Exploratory moderator analyses (random-effects categorical or metaregression models in R using the metafor package) explored potential explanations for the between-subject variance of effect sizes. These included 1) drug type (agonist or antagonist), 2) more specific dopamine drug activity types (D2 antagonist, D2 agonist, dopamine precursor or dopamine reuptake inhibitor), 3) task type, 4) dose regimen (single or multiple), 5) mean age across the study and 6) the mean proportion of female participants included in the study. The $R^2$ value was used to quantify the proportion of between-study heterogeneity accounted for by the moderators.

##### 1.3.5 Assessment of study quality

The quality of studies were assessed with the Cochrane Collaboration's tool for assessing risk of bias[58] independently by Z.Q., Q.D., and T.S.

Assessed criteria:
1. Random sequence generation (selection bias)
2. Allocation concealment (selection bias)
3. Blinding of participants and researchers (performance bias)
4. Blinding of outcome assessment (detection bias)
5. Incomplete outcome data (attrition bias)

For each item, the risk of bias was marked as 'low' if it was explicitly mentioned, 'high' if it was not explicitly mentioned, and 'unclear' if it was not explicitly mentioned but might be true.

#### 1.4 Effect size calculations

The SMDs for between-subject studies were calculated using Cohen's *d*[59,60]. These were extracted from papers reporting the required statistics or by using a plot digitizer (PlotDigitizer, 3.1.6, 2024). Both within- and between-subject designs were included since this design choice is irrelevant to the question and there were a substantial number of each (23 between-subject dopamine studies, 45 within-subject dopamine studies; 25 between-subject serotonin studies, 14 within-subject studies).

However, combining within- and between-subject data in one meta-analysis requires knowledge of the correlation between the drug and placebo condition from within-subject studies in order to calculate an equivalent (correlation-corrected) effect size to the between-subject design Cohen's *d*[61,62]. Since none of the included studies reported this, we only included within-subject studies where we could either derive the correlation from the information provided (i.e., where the mean and SD of the drug and placebo condition, and a t-statistic or standard error of the difference were reported) or data were provided from contacted authors. For these within-subject studies, we calculated the Cohen's $d_{rm}$ and associated variance using the Cousineau and Goulet-Pelletier [61] method in the TOSTER package[63]. We also compared this to alternatives in both the effect size and variance computation for within-subject studies, which made little difference (eFigure 1). If a within-subject study only provided enough data for calculating between-subject Cohen's *d* and raw data was unavailable, we classified it as a between-subject study.

Some within-subject studies applied mixed-effects models (due to uneven subjects in each condition) or non-parametric methods to analyze the data. Since calculating Cohen's $d_{rm}$ is not straightforward in these cases, we opted to treat these as between-subject studies if the sample size in each group was very low for a within-subject comparison or transform the raw data, respectively.

In total, out of the original within-subject studies included, 13 studies in the dopamine domain and 4 studies in the serotonin domain were treated as between-subject studies. For meta-analyses where we additionally analyzed subcomponents based on only within-subject studies, we computed Cohen's $d_z$. These included 3 additional serotonin and 5 additional dopamine studies.

##### 1.4.1 Conversion of outcome measures into effect sizes and associated variances

For between-subject studies and studies treated as between-subject studies, the following equations were used:

###### *1.4.1.1 Cohen's d from Means and Standard Deviations of 2 samples*

$$d = \frac{M_1 - M_2}{SD_{pooled}}, \tag{1}$$

where *d* is Cohen's *d*, $M_1$ is the mean of one sample, $M_2$ is the mean of the other sample, $SD_{pooled}$ is the pooled standard deviation of the two samples.

###### *1.4.1.2 Pooled standard deviation of 2 samples*

$$SD_{pooled} = \sqrt{\frac{(N_1 - 1)SD_1^2 + (N_2 - 1)SD_2^2}{N_1 + N_2 - 2}}, \tag{2}$$

where $N_1$ is the size of one sample, $N_2$ is the size of the other sample, $SD_1$ is the standard deviation of one sample, $SD_2$ is the standard deviation of the other sample.

###### *1.4.1.3 Cohen's d from t-statistic*

$$d = \frac{t}{\sqrt{\frac{1}{N_1} + \frac{1}{N_2}}}, \tag{3}$$

where *t* is the independent t-statistic reported.

###### *1.4.1.4 Cohen's d from F-statistic*

$$SD_{pooled} = \sqrt{\frac{F}{\frac{1}{N_1} + \frac{1}{2}}}, \tag{4}$$

where *F* is the F-statistic reported.

###### *1.4.1.5 Variance on Cohen's d*

$$var(d) = \frac{N_1 + N_2}{N_1 \times N_2} + \frac{d^2}{2(N_1 + N_2)} \tag{5}$$

For within-subject studies, Cohen's $d_{rm}$ was computed, taking into account the correlation between measures, using the Cousineau and Goulet-Pelletier [61] method. This is useful for when effect sizes are being compared for studies that involve between and within subjects designs[60]. The TOSTER package in R was used, which uses the following equations[63]:

##### *1.4.1.6 Cohen's $d_{rm}$ from Means, Standard Deviations and Correlation Coefficient of paired samples*

$$d_{rm} = \frac{M_1 - M_2}{S_{diff}} \times \sqrt{\frac{2}{(1-r)}} \tag{6}$$

where $d_{rm}$ is Cohen's $d_{rm}$, $M_1$ is the mean of one condition (drug), $M_2$ is the mean of the other condition (placebo), $S_{diff}$ is the standard deviation of the difference scores, and $r$ is the correlation between the drug and placebo conditions.

##### *1.4.1.7 The standard deviation of the difference scores*

$$S_{diff} = \sqrt{SD_1^2 + SD_2^2 - 2 \times r \times SD_1 \times SD_2}, \tag{7}$$

where $S_{diff}$ is the standard deviation of the difference scores of the two samples, $SD_1$ is the standard deviation of one condition, and $SD_2$ is the standard deviation of the other condition.

##### *1.4.1.8 The degrees of freedom for Cohen's $d_{rm}$ under the Cousineau and Goulet-Pelletier (2021) method*

$$df = 2(N-1), \tag{8}$$

where $N$ represents the sample size.

##### *1.4.1.9 The variance of Cohen's $d_{rm}$*

$$var(d_{rm}) = \frac{df}{df-2} \times \frac{2(1-r)}{N} \times (1 + d_{rm}^2 \times \frac{N}{2(1-r)}) - \frac{d_{rm}^2}{1} \tag{9}$$

For within-subject studies where we had $M_1$, $M_2$, $SD_1$, $SD_2$ and the paired t-statistic, we estimated the correlation coefficient $r$ between conditions with the following procedure.

Compute standard error of the difference, *SED*:

$$SED = \left|\frac{M_1 - M_2}{t}\right| \quad (10)$$

Compute the standard deviation of the difference scores, $S_{diff}$:

$$S_{diff} = SED \times \sqrt{N} \quad (11)$$

Calculate covariance, *Cov*:

$$Cov = \frac{SD_1^2 + SD_2^2 - S_{diff}^2}{2} \quad (12)$$

The correlation coefficient *r* is:

$$r = \frac{Cov}{SD_1 \times SD_2} \quad (13)$$

For supplemental meta-analyses where we only included within-subject studies, we first computed a paired t-test if one was not already provided in the study and then computed Cohen's $d_z$ with the following equation:

*1.4.1.10 Cohen's $d_z$ based on the t-statistic, t*

$$d_z = \frac{t}{\sqrt{N}} \quad (14)$$

*1.4.1.11 The variance of Cohen's $d_z$*

$$var(d_z) = \frac{1}{N} + \frac{d_z^2}{2 \times N} \quad (15)$$

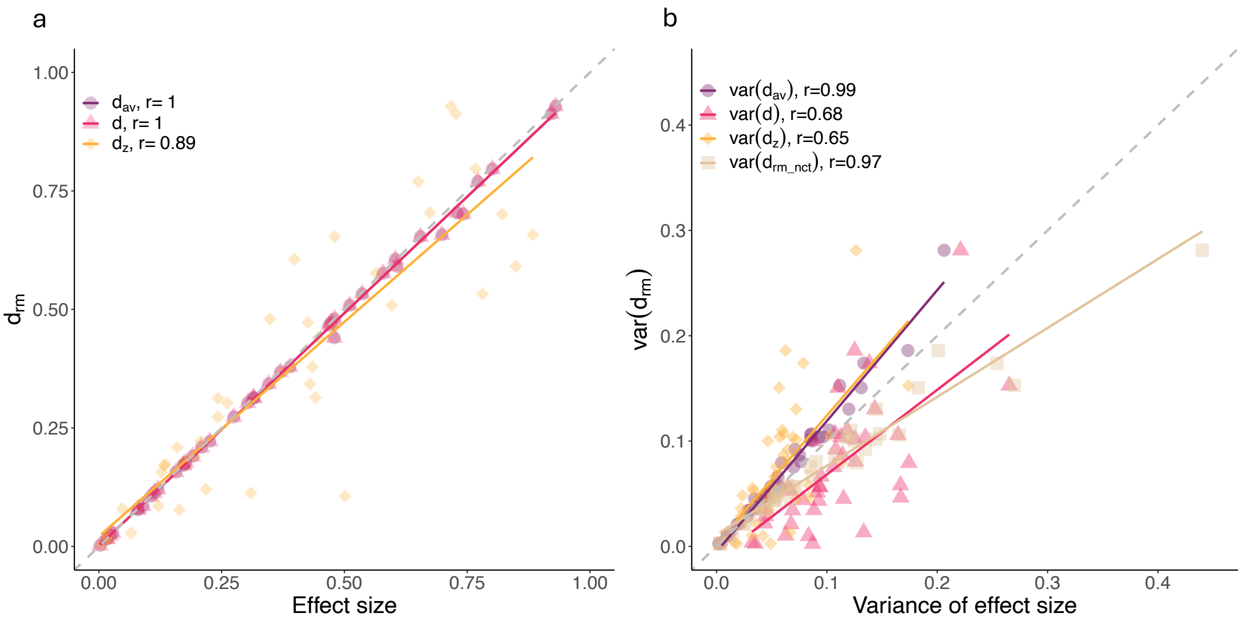


**eFigure 1: Comparison of a) effect size and b) variance of effect size calculations for within-subject studies that provided sufficient data to compute all variations (N=42).** The y-axis represents Cohen's $d_{rm}$ and the x-axis represents Cohen's $d_{av}$, $d$, or $d_z$ in figure a. $d_{av} = \frac{M_1 - M_2}{\sqrt{\frac{SD_1^2 + SD_2^2}{2}}}$[59], and will equal $d$ when the condition sample sizes are equal. In figure b, the y-axis represents the variance of Cohen's $d_{rm}$ (equation 9) and the x-axis represents four other calculations of the variance. *Var(d)* represents the variance of Cohen's $d$ (equation 5); *var*($d_z$) represents the variance of $d_z$ (equation 15); and *var*($d_{av}$) represents the variance of $d_{av}$. *Var*($d_{av}$) is calculated as: $var(d_{av}) = (\frac{CI_{up} - CI_{low}}{2\times 1.96})^2$, where CI is calculated using the non-central $t$ distribution: $CI = d_{av} \pm t(1 - \frac{\alpha}{2}, n - 1, \lambda) \times \sqrt{\frac{2(SD_1^2 + SD_2^2 - 2\times r\times SD_1 \times SD_2)}{N(SD_1^2 + SD_2^2)}}$, $\alpha$=0.05 and the non-centrality parameter is the observed t-value: $\lambda = d_{av} \times \sqrt{\frac{N(SD_1^2 + SD_2^2)}{2(SD_1^2 + SD_2^2 - 2\times r\times SD_1 \times SD_2)}}$[64]. *Var*($d_{rm\_nct}$) represents the variance of $d_{rm}$ calculated with the setting "nct" for the confidence interval type in the package TOSTER[63], which uses the same equation as in equation 9 but with $df = N - 1$. The gray dished diagonal line represents the reference line when y=x.

### 2 Supplementary results

| | Dopamine | | | Serotonin | | |
|---|---|---|---|---|---|---|
| **Category** | **No. of studies** | **Placebo N** | **Drug N** | **No. of studies** | **Placebo N** | **Drug N** |
| Overall reward | 49 | 1260 | 1266 | 21 | 572 | 559 |
| Overall punishment | 17 | 362 | 365 | 11 | 292 | 285 |
| Reward learning/sensitivity | 25 | 616 | 623 | 9 | 274 | 269 |
| Reward discounting | 12 | 439 | 436 | 4 | 74 | 74 |
| Reward response vigour | 9 | 185 | 186 | 4 | 96 | 94 |
| Appetitive Pavlovian | 5 | 158 | 157 | 4 | 128 | 122 |
| Punishment learning/sensitivity | 13 | 244 | 249 | 4 | 108 | 106 |
| Aversive Pavlovian | 4 | 118 | 116 | 7 | 184 | 179 |
| Model-based learning/flexibility | 8 | 236 | 239 | 10 | 306 | 300 |
| Risk attitude | 5 | 155 | 152 | 6 | 105 | 104 |
| Aversive Pavlovian within-subjects | - | - | - | 6 | 145 | - |
| Risk attitude within-subjects | 5 | 127 | - | - | - | - |
| Model-based learning/flexibility within-subjects | 8 | 198 | - | - | - | - |

**eTable 2: Number of studies and participants (N) for each condition for each assessed category.**

### 2.1 Assessment of study quality

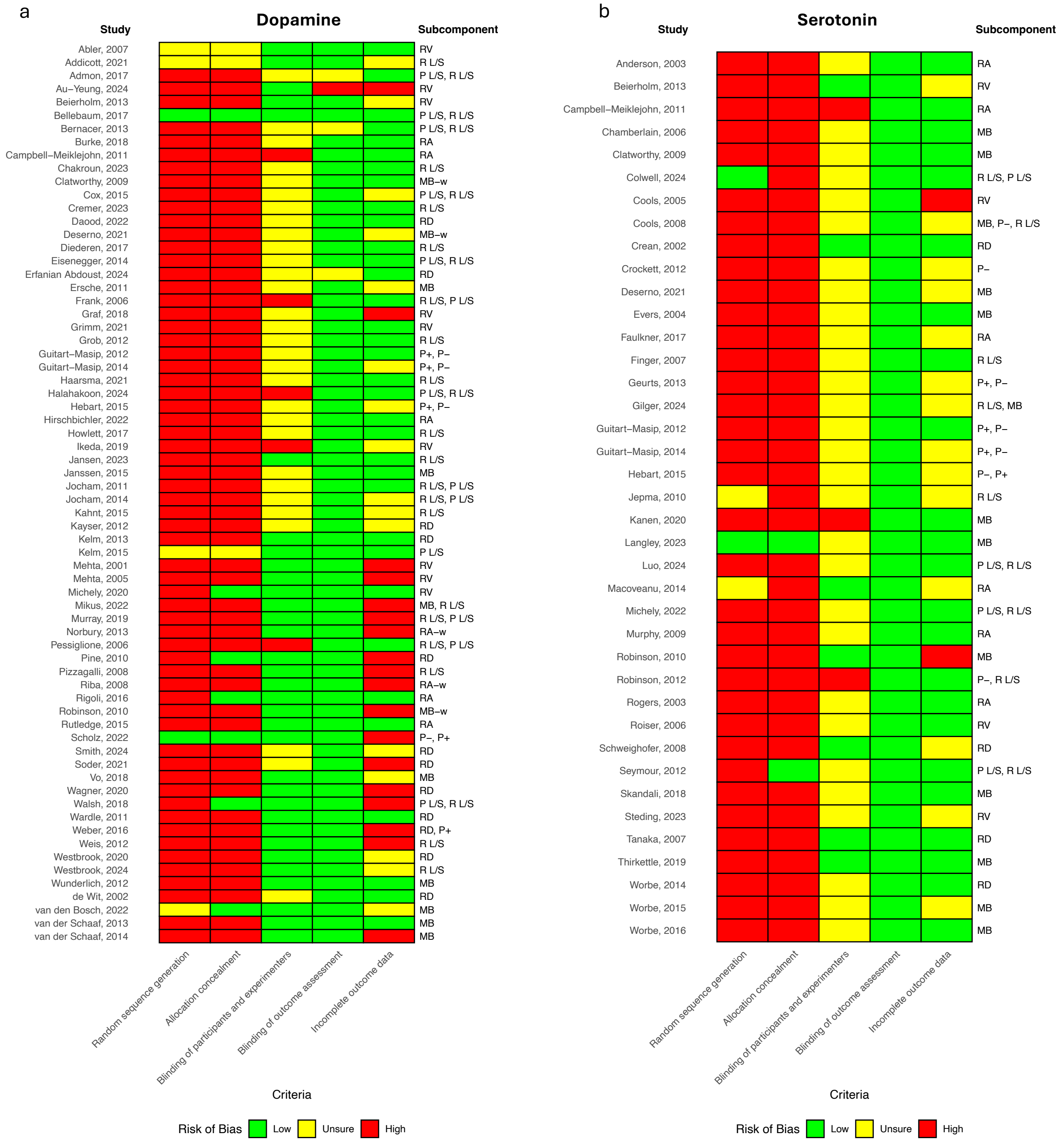


**eFigure 2: Risk of bias assessment for a) dopamine and b) serotonin studies.** Abbreviations: R L/S: reward learning sensitivity; P L/S: punishment learning/sensitivity; P+: appetitive Pavlovian; P-: aversive Pavlovian; RD: reward discounting; RV: reward vigor; MB: model-based learning/flexibility; RA: risk attitude: -w: within-subjects study only

#### 2.2 Additional dopamine results

a

| Study | N(Drug, Pla) | Drug, dose | SMD [95% CI] |
|---|---|---|---|
| Jocham, 2011 | 16 | Amisulpride 200mg | 0.93 [ 0.11, 1.75] * |
| Admon, 2017 | 20, 18 | Amisulpride 50mg | 0.26 [−0.38, 0.90] * |
| Frank, 2006 | 19, 20 | Haloperidol 2mg | 0.37 [−0.26, 1.00] * |
| Wagner, 2020 | 23, 26 | Haloperidol 2mg | 0.53 [−0.04, 1.10] * |
| Erfanian Abdoust, 2024 | 62 | Haloperidol 2mg | 0.00 [−0.12, 0.12] * |
| Walsh, 2018 | 20, 20 | Bupropion 150mg | −0.06 [−0.68, 0.56] |
| Ikeda, 2019 | 15 | Bupropion 150mg | 0.30 [−0.33, 0.93] |
| de Wit, 2002 | 36, 36 | D−amphetamine 20mg | 0.21 [−0.25, 0.67] |
| Wardle, 2011 | 17, 17 | D−amphetamine 20mg | 0.64 [−0.05, 1.33] |
| Soder, 2021 | 28, 28 | D−amphetamine 20mg | 0.33 [−0.20, 0.86] |
| Bernacer, 2013 | 17 | Methamphetamine 0.3mg/kg | −0.70 [−1.33, −0.08] |
| Westbrook, 2020 | 46 | Methylphenidate 20mg | 0.19 [−0.10, 0.48] |
| Daood, 2022 | 57 | Methylphenidate 20mg | 0.11 [ 0.01, 0.21] |
| Howlett, 2017 | 19 | Methylphenidate 40mg | 0.27 [−0.26, 0.81] |
| Addicott, 2021 | 22 | Methylphenidate 40mg | 0.38 [−0.03, 0.79] |
| Bellebaum, 2017 | 16, 20 | Modafinil 200mg | 0.38 [−0.29, 1.04] |
| Pessiglione, 2006 | 13, 13 | L−dopa 100mg+25mg | 0.75 [−0.04, 1.55] |
| Grimm, 2021 | 45 | L−dopa 100mg+25mg | 0.02 [−0.35, 0.39] |
| Jansen, 2023 | 30, 30 | L−dopa 100mg+25mg | 0.00 [−0.51, 0.51] |
| Weis, 2012 | 27, 28 | L−dopa 100mg+35mg | 0.04 [−0.49, 0.57] |
| Pine, 2010 | 13 | L−dopa 150mg | −0.70 [−1.34, −0.06] |
| Michely, 2020 | 20 | L−dopa 150mg | 0.80 [ 0.20, 1.39] |
| Smith, 2024 | 76, 76 | L−dopa 150mg | 0.14 [−0.18, 0.46] |
| Guitart−Masip, 2012 | 16, 20 | L−dopa 150mg+37.5mg | −0.10 [−0.76, 0.56] |
| Beierholm, 2013 | 30, 30 | L−dopa 150mg+37.5mg | 0.59 [ 0.07, 1.11] |
| Guitart−Masip, 2014 | 30, 29 | L−dopa 150mg+37.5mg | −0.30 [−0.82, 0.21] |
| Chakroun, 2023 | 31 | L−dopa 150mg+37.5mg | −0.17 [−0.63, 0.29] |
| Mehta, 2001 | 18 | Bromocriptine 1.25mg | 0.47 [−0.10, 1.05] |
| Au−Yeung, 2024 | 19, 18 | Pramipexole 1mg/day | 0.21 [−0.44, 0.86] |
| Halahakoon, 2024 | 21, 19 | Pramipexole 1mg/day | 0.68 [ 0.04, 1.32] |
| Kayser, 2012 | 23 | Tolcapone 200mg | 0.11 [ 0.01, 0.20] |
| Scholz, 2022 | 35, 35 | Tolcapone 200mg | 0.03 [−0.44, 0.50] |
| Graf, 2018 | 17 | Amisulpride 200mg/day | 0.21 [−0.44, 0.86] |
| Jocham, 2014 | 22 | Amisulpride 400mg | 0.44 [−0.02, 0.90] |
| Kahnt, 2015 | 27, 24 | Amisulpride 400mg | 0.24 [−0.31, 0.79] |
| Weber, 2016 | 40, 40 | Amisulpride 400mg | −0.58 [−1.02, −0.13] |
| Murray, 2019 | 18, 18 | Amisulpride 400mg | −0.04 [−0.69, 0.62] |
| Mikus, 2022 | 38, 35 | Amisulpride 400mg | 0.38 [−0.08, 0.84] |
| Cremer, 2023 | 23, 22 | Amisulpride 400mg | 0.59 [−0.00, 1.19] |
| Abler, 2007 | 8 | Olanzapine 5mg | 0.66 [−0.11, 1.42] |
| Diederen, 2017 | 19, 19 | Sulpiride 600mg | 0.58 [−0.07, 1.23] |
| Haarsma, 2021 | 20, 20 | Sulpiride 600mg | 0.71 [ 0.07, 1.35] |
| Eisenegger, 2014 | 41, 35 | Sulpiride 800mg | −0.05 [−0.50, 0.40] |
| Grob, 2012 | 28, 28 | AMPT 40mg/kg | −0.36 [−0.88, 0.17] |
| Cox, 2015 | 10, 10 | Phe / Tyr depletion 100g | 0.00 [−0.88, 0.88] |
| Mehta, 2005 | 14 | Phe / Tyr depletion | −0.17 [−0.88, 0.54] |
| Hebart, 2015 | 35, 34 | Phe / Tyr depletion | 0.65 [ 0.17, 1.14] |
| Kelm, 2013 | 15 | Phe / Tyr depletion 68.7g | −0.03 [−0.26, 0.20] |
| Pizzagalli, 2008 | 11, 13 | Pramipexole 0.5mg | 1.42 [ 0.52, 2.32] * |
| Random−effects meta−analysis: | | | 0.18 [ 0.09, 0.28] |

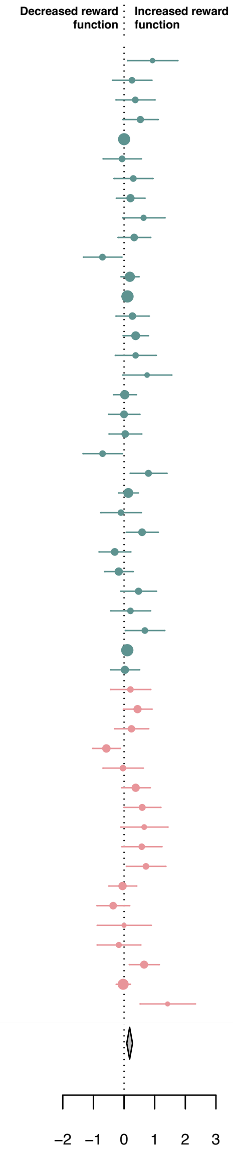


b

| Study | N(Drug, Pla) | Drug, dose | SMD [95% CI] |
|---|---|---|---|
| Jocham, 2011 | 16 | Amisulpride 200mg | 0.08 [−0.77, 0.92]* |
| Admon, 2017 | 20, 18 | Amisulpride 50mg | 0.54 [−0.11, 1.19]* |
| Frank, 2006 | 19, 20 | Haloperidol 2mg | −0.38 [−1.01, 0.25]* |
| Walsh, 2018 | 20, 20 | Bupropion 150mg | 0.37 [−0.25, 1.00] |
| Bernacer, 2013 | 17, 17 | Methamphetamine 0.3mg/kg | −0.01 [−0.68, 0.66] |
| Bellebaum, 2017 | 16, 20 | Modafinil 200mg | −0.52 [−1.19, 0.14] |
| Pessiglione, 2006 | 13, 13 | L−dopa 100mg+25mg | 0.42 [−0.35, 1.20] |
| Guitart−Masip, 2012 | 16, 20 | L−dopa 150mg+37.5mg | 0.19 [−0.47, 0.85] |
| Guitart−Masip, 2014 | 30, 29 | L−dopa 150mg+37.5mg | −0.21 [−0.72, 0.30] |
| Halahakoon, 2024 | 21, 19 | Pramipexole 1mg/day | −0.63 [−1.27, 0.00] |
| Scholz, 2022 | 35, 35 | Tolcapone 200mg | −0.09 [−0.56, 0.37] |
| Jocham, 2014 | 22 | Amisulpride 400mg | 0.58 [ 0.07, 1.08] |
| Murray, 2019 | 18, 18 | Amisulpride 400mg | −0.51 [−1.17, 0.16] |
| Eisenegger, 2014 | 41, 35 | Sulpiride 800mg | 0.22 [−0.23, 0.67] |
| Cox, 2015 | 10 | Phe / Tyr depletion 100g | −0.91 [−1.95, 0.13] |
| Hebart, 2015 | 35, 34 | Phe / Tyr depletion | −0.40 [−0.88, 0.07] |
| Kelm, 2015 | 16 | Phe / Tyr depletion 68.7g | −0.22 [−0.78, 0.33] |
| Random−effects meta−analysis: | | | −0.06 [−0.26, 0.13] |

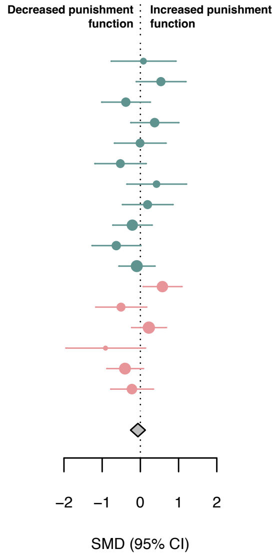


**eFigure 3: The effect of upregulating dopamine on overall reward and punishment.** Standardized mean differences (SMDs) between the effect of upregulating dopamine versus placebo on a) overall reward function, and b) overall punishment function. Pla indicates placebo. We recoded antagonist effects as if they were agonists (pink SMD point estimates and 95% CI). Green SMD point estimates and 95% CI indicate studies that were coded as original agonists. Asterisks indicate studies that used a low dose of an agonist or antagonists. These effects were interpreted contrary to their typical activity profile (e.g., a low-dose agonist acting antagonistically), as suggested by the literature.

a

| Study | N(Drug, Pla) | Drug, dose | SMD [95% CI] |
|---|---|---|---|
| Walsh, 2018 | 20, 20 | Bupropion 150mg | −0.06 [−0.68, 0.56] |
| Ikeda, 2019 | 15 | Bupropion 150mg | 0.30 [−0.33, 0.93] |
| de Wit, 2002 | 36, 36 | D−amphetamine 20mg | 0.21 [−0.25, 0.67] |
| Wardle, 2011 | 17, 17 | D−amphetamine 20mg | 0.64 [−0.05, 1.33] |
| Soder, 2021 | 28, 28 | D−amphetamine 20mg | 0.33 [−0.20, 0.86] |
| Bernacer, 2013 | 17 | Methamphetamine 0.3mg/kg | −0.70 [−1.33, −0.08] |
| Westbrook, 2020 | 46 | Methylphenidate 20mg | 0.19 [−0.10, 0.48] |
| Daood, 2022 | 57 | Methylphenidate 20mg | 0.11 [ 0.01, 0.21] |
| Howlett, 2017 | 19 | Methylphenidate 40mg | 0.27 [−0.26, 0.81] |
| Addicott, 2021 | 22 | Methylphenidate 40mg | 0.38 [−0.03, 0.79] |
| Bellebaum, 2017 | 16, 20 | Modafinil 200mg | 0.38 [−0.29, 1.04] |
| Pessiglione, 2006 | 13, 13 | L−dopa 100mg+25mg | 0.75 [−0.04, 1.55] |
| Grimm, 2021 | 45 | L−dopa 100mg+25mg | 0.02 [−0.35, 0.39] |
| Jansen, 2023 | 30, 30 | L−dopa 100mg+25mg | 0.00 [−0.51, 0.51] |
| Weis, 2012 | 27, 28 | L−dopa 100mg+35mg | 0.04 [−0.49, 0.57] |
| Pine, 2010 | 13 | L−dopa 150mg | −0.70 [−1.34, −0.06] |
| Michely, 2020 | 20 | L−dopa 150mg | 0.80 [ 0.20, 1.39] |
| Smith, 2024 | 76, 76 | L−dopa 150mg | 0.14 [−0.18, 0.46] |
| Guitart−Masip, 2012 | 16, 20 | L−dopa 150mg+37.5mg | −0.10 [−0.76, 0.56] |
| Beierholm, 2013 | 30, 30 | L−dopa 150mg+37.5mg | 0.59 [ 0.07, 1.11] |
| Guitart−Masip, 2014 | 30, 29 | L−dopa 150mg+37.5mg | −0.30 [−0.82, 0.21] |
| Chakroun, 2023 | 31 | L−dopa 150mg+37.5mg | −0.17 [−0.63, 0.29] |
| Mehta, 2001 | 18 | Bromocriptine 1.25mg | 0.47 [−0.10, 1.05] |
| Pizzagalli, 2008 | 11, 13 | Pramipexole 0.5mg | −1.42 [−2.32, −0.52] |
| Au−Yeung, 2024 | 19, 18 | Pramipexole 1mg/day | 0.21 [−0.44, 0.86] |
| Halahakoon, 2024 | 21, 19 | Pramipexole 1mg/day | 0.68 [ 0.04, 1.32] |
| Kayser, 2012 | 23 | Tolcapone 200mg | 0.11 [ 0.01, 0.20] |
| Scholz, 2022 | 35, 35 | Tolcapone 200mg | 0.03 [−0.44, 0.50] |
| Jocham, 2011 | 16 | Amisulpride 200mg | −0.93 [−1.75, −0.11] |
| Graf, 2018 | 17 | Amisulpride 200mg/day | 0.21 [−0.44, 0.86] |
| Jocham, 2014 | 22 | Amisulpride 400mg | 0.44 [−0.02, 0.90] |
| Kahnt, 2015 | 27, 24 | Amisulpride 400mg | 0.24 [−0.31, 0.79] |
| Weber, 2016 | 40, 40 | Amisulpride 400mg | −0.58 [−1.02, −0.13] |
| Murray, 2019 | 18, 18 | Amisulpride 400mg | −0.04 [−0.69, 0.62] |
| Mikus, 2022 | 38, 35 | Amisulpride 400mg | 0.38 [−0.08, 0.84] |
| Cremer, 2023 | 23, 22 | Amisulpride 400mg | 0.59 [−0.00, 1.19] |
| Admon, 2017 | 20, 18 | Amisulpride 50mg | −0.26 [−0.90, 0.38] |
| Frank, 2006 | 19, 20 | Haloperidol 2mg | −0.37 [−1.00, 0.26] |
| Wagner, 2020 | 23, 26 | Haloperidol 2mg | −0.53 [−1.10, 0.04] |
| Erfanian Abdoust, 2024 | 62 | Haloperidol 2mg | −0.00 [−0.12, 0.12] |
| Abler, 2007 | 8 | Olanzapine 5mg | 0.66 [−0.11, 1.42] |
| Diederen, 2017 | 19, 19 | Sulpiride 600mg | 0.58 [−0.07, 1.23] |
| Haarsma, 2021 | 20, 20 | Sulpiride 600mg | 0.71 [ 0.07, 1.35] |
| Eisenegger, 2014 | 41, 35 | Sulpiride 800mg | −0.05 [−0.50, 0.40] |
| Grob, 2012 | 28, 28 | AMPT 40mg/kg | −0.36 [−0.88, 0.17] |
| Cox, 2015 | 10, 10 | Phe / Tyr depletion 100g | 0.00 [−0.88, 0.88] |
| Mehta, 2005 | 14 | Phe / Tyr depletion | −0.17 [−0.88, 0.54] |
| Hebart, 2015 | 35, 34 | Phe / Tyr depletion | 0.65 [ 0.17, 1.14] |
| Kelm, 2013 | 15 | Phe / Tyr depletion 68.7g | −0.03 [−0.26, 0.20] |
| Random−effects meta−analysis: | | | 0.10 [−0.00, 0.20] |

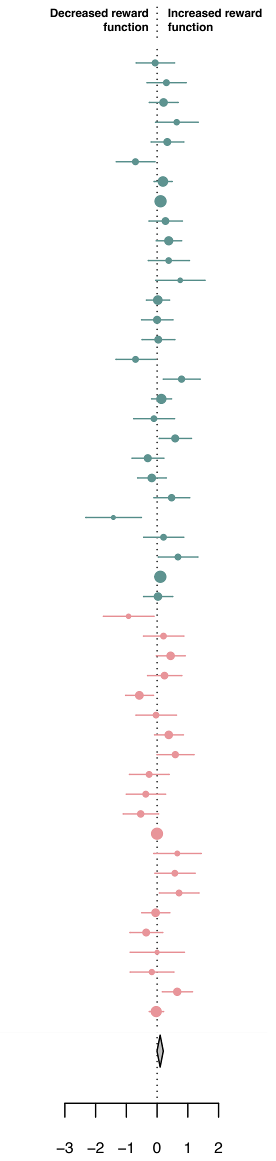


b

| Study | N(Drug, Pla) | Drug, dose | SMD [95% CI] |
|---|---|---|---|
| Walsh, 2018 | 20, 20 | Bupropion 150mg | 0.37 [−0.25, 1.00] |
| Bernacer, 2013 | 17, 17 | Methamphetamine 0.3mg/kg | −0.01 [−0.68, 0.66] |
| Bellebaum, 2017 | 16, 20 | Modafinil 200mg | −0.52 [−1.19, 0.14] |
| Pessiglione, 2006 | 13, 13 | L−dopa 100mg+25mg | 0.42 [−0.35, 1.20] |
| Guitart−Masip, 2012 | 16, 20 | L−dopa 150mg+37.5mg | 0.19 [−0.47, 0.85] |
| Guitart−Masip, 2014 | 30, 29 | L−dopa 150mg+37.5mg | −0.21 [−0.72, 0.30] |
| Halahakoon, 2024 | 21, 19 | Pramipexole 1mg/day | −0.63 [−1.27, 0.00] |
| Scholz, 2022 | 35, 35 | Tolcapone 200mg | −0.09 [−0.56, 0.37] |
| Jocham, 2011 | 16 | Amisulpride 200mg | −0.08 [−0.92, 0.77] |
| Jocham, 2014 | 22 | Amisulpride 400mg | 0.58 [ 0.07, 1.08] |
| Murray, 2019 | 18, 18 | Amisulpride 400mg | −0.51 [−1.17, 0.16] |
| Admon, 2017 | 20, 18 | Amisulpride 50mg | −0.54 [−1.19, 0.11] |
| Frank, 2006 | 19, 20 | Haloperidol 2mg | 0.38 [−0.25, 1.01] |
| Eisenegger, 2014 | 41, 35 | Sulpiride 800mg | 0.22 [−0.23, 0.67] |
| Cox, 2015 | 10 | Phe / Tyr depletion 100g | −0.91 [−1.95, 0.13] |
| Hebart, 2015 | 35, 34 | Phe / Tyr depletion | −0.40 [−0.88, 0.07] |
| Kelm, 2015 | 16 | Phe / Tyr depletion 68.7g | −0.22 [−0.78, 0.33] |
| Random−effects meta−analysis: | | | −0.09 [−0.28, 0.11] |

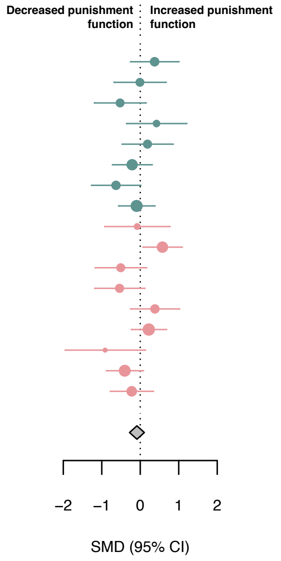


**eFigure 4: The effect of upregulating dopamine on overall reward and punishment according to original drug profile.** Standardized mean differences (SMDs) between the effect of upregulating dopamine versus placebo on a) overall reward function, and b) overall punishment function according to the original drug activity profile. Pla indicates placebo. We recoded antagonist effects as if they were agonists (pink SMD point estimates and 95% CI). Green SMD point estimates and 95% CI indicate studies that were coded as original agonists.

a

| Study | N(Drug, Pla) | Drug, dose | SMD [95% CI] |
|---|---|---|---|
| Guitart–Masip, 2012 | 16, 20 | L–dopa 150mg+37.5mg | −0.10 [−0.76, 0.56] |
| Guitart–Masip, 2014 | 30, 29 | L–dopa 150mg+37.5mg | −0.30 [−0.82, 0.21] |
| Scholz, 2022 | 35, 35 | Tolcapone 200mg | 0.03 [−0.44, 0.50] |
| Weber, 2016 | 41, 40 | Amisulpride 400mg | 0.67 [ 0.22, 1.12] |
| Hebart, 2015 | 35, 34 | Phe / Tyr depletion | 0.65 [ 0.17, 1.14] |
| Random–effects meta–analysis: | | | 0.21 [−0.19, 0.61] |

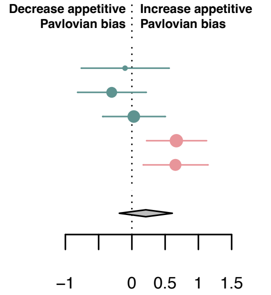


b

| Study | N(Drug, Pla) | Drug, dose | SMD [95% CI] |
|---|---|---|---|
| Hirschbichler, 2022 | 30 | L–dopa 100mg | 0.51 [ 0.15, 0.87] |
| Rutledge, 2015 | 30 | L–dopa 150mg+37.5mg | 0.31 [ 0.03, 0.60] |
| Rigoli, 2016 | 32 | L–dopa 150mg+37.5mg | 0.12 [−0.08, 0.32] |
| Burke, 2018 | 45, 48 | Amisulpride 400mg | −0.79 [−1.21, −0.37] |
| Campbell–Meiklejohn, 2011 | 15, 15 | Pramipexole 0.176μg | 0.31 [−0.41, 1.03] * |
| Random–effects meta–analysis: | | | 0.09 [−0.36, 0.54] |

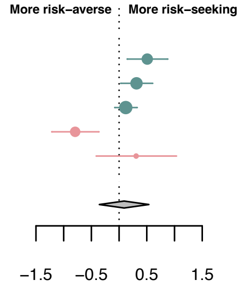


c

| Study | N(Drug, Pla) | Drug, dose | SMD [95% CI] |
|---|---|---|---|
| van der Schaaf, 2013 | 19 | Methylphenidate 20mg | 0.61 [−0.16, 1.37] |
| van den Bosch, 2022 | 88, 88 | Methylphenidate 20mg | 0.45 [ 0.15, 0.75] |
| Vo, 2018 | 13, 13 | L–dopa 100mg+25mg | −1.12 [−1.94, −0.29] |
| Wunderlich, 2012 | 18 | L–dopa 150mg+37.5mg | 0.53 [ 0.12, 0.95] |
| Ersche, 2011 | 18 | Amisulpride 400mg | 0.31 [−0.32, 0.94] |
| Mikus, 2022 | 38, 35 | Amisulpride 400mg | −0.76 [−1.23, −0.28] |
| van der Schaaf, 2014 | 23 | Sulpiride 400mg | −0.34 [−0.71, 0.02] |
| Janssen, 2015 | 22 | Sulpiride 400mg | 0.47 [−0.00, 0.93] |
| Random–effects meta–analysis: | | | 0.04 [−0.39, 0.47] |

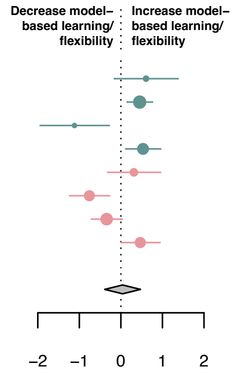


d

| Study | N(Drug, Pla) | Drug, dose | SMD [95% CI] |
|---|---|---|---|
| Norbury, 2013 | 20 | Cabergoline 1.5mg | −0.16 [−0.60, 0.28] |
| Hirschbichler, 2022 | 30 | L–dopa 100mg | 0.60 [ 0.21, 0.99] |
| Rutledge, 2015 | 30 | L–dopa 150mg+37.5mg | 0.44 [ 0.07, 0.82] |
| Rigoli, 2016 | 32 | L–dopa 150mg+37.5mg | 0.22 [−0.13, 0.57] |
| Riba, 2008 | 15 | Pramipexole 0.5mg | −0.26 [−0.77, 0.26] * |
| Random–effects meta–analysis: | | | 0.19 [−0.12, 0.50] |

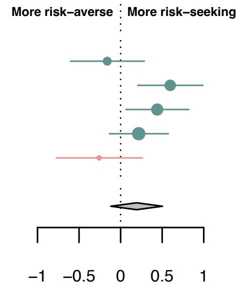


e

| Study | N(Drug, Pla) | Drug, dose | SMD [95% CI] |
|---|---|---|---|
| van der Schaaf, 2013 | 19 | Methylphenidate 20mg | 0.40 [−0.07, 0.87] |
| Clatworthy, 2009 | 9 | Methylphenidate 60mg | −0.04 [−0.69, 0.61] |
| Wunderlich, 2012 | 18 | L–dopa 150mg+37.5mg | 0.78 [ 0.25, 1.31] |
| Deserno, 2021 | 62 | L–dopa 150mg+37.5mg | −0.05 [−0.30, 0.20] |
| Robinson, 2010 | 27 | Phe/Tyr depletion | −0.18 [−0.56, 0.20] |
| Ersche, 2011 | 18 | Amisulpride 400mg | 0.24 [−0.23, 0.71] |
| van der Schaaf, 2014 | 23 | Sulpiride 400mg | −0.43 [−0.86, −0.00] |
| Janssen, 2015 | 22 | Sulpiride 400mg | 0.48 [ 0.03, 0.92] |
| Random–effects meta–analysis: | | | 0.13 [−0.14, 0.40] |

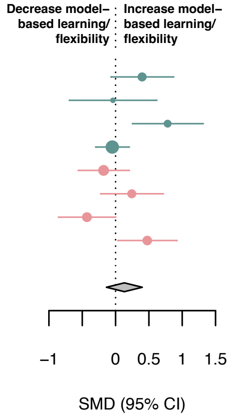


**eFigure 5: The effect of upregulating dopamine on appetitive Pavlovian bias, risk and model-based learning/flexibility.** Standardized mean differences (SMDs) between the effect of upregulating dopamine versus placebo on a) appetitive Pavlovian bias, b) risk attitude, c) model-based learning/flexibility, d) risk attitude in within-subject studies only, and e) model-based learning/flexibility in within-subjects only. Forest plots d and e display SMDs that reflect Cohen's $d_z$. Pla indicates placebo. We recoded antagonist effects as if they were agonists (pink SMD point estimates and 95% CI). Green SMD point estimates and 95% CI indicate studies that were coded as original agonists. Asterisks indicate studies that used a low dose of an agonist or antagonists. These effects were interpreted contrary to their typical activity profile (e.g., a low-dose agonist acting antagonistically), as suggested by the literature.

a

| Study | N(Drug, Pla) | Drug, dose | SMD [95% CI] |
|---|---|---|---|
| Walsh, 2018 | 20, 20 | Bupropion 150mg | −0.06 [−0.68, 0.56] |
| Bernacer, 2013 | 17 | Methamphetamine 0.3mg/kg | −0.70 [−1.33, −0.08] |
| Westbrook, 2024 | 95, 97 | Methylphenidate 20mg | 0.37 [ 0.09, 0.66] |
| Howlett, 2017 | 19 | Methylphenidate 40mg | 0.27 [−0.26, 0.81] |
| Addicott, 2021 | 22 | Methylphenidate 40mg | 0.38 [−0.03, 0.79] |
| Bellebaum, 2017 | 16, 20 | Modafinil 200mg | 0.38 [−0.29, 1.04] |
| Pessiglione, 2006 | 13, 13 | L−dopa 100mg+25mg | 0.75 [−0.04, 1.55] |
| Jansen, 2023 | 30, 30 | L−dopa 100mg+25mg | 0.00 [−0.51, 0.51] |
| Weis, 2012 | 27, 28 | L−dopa 100mg+35mg | 0.04 [−0.49, 0.57] |
| Chakroun, 2023 | 31 | L−dopa 150mg+37.5mg | −0.17 [−0.63, 0.29] |
| Pizzagalli, 2008 | 11, 13 | Pramipexole 0.5mg | −1.42 [−2.32, −0.52] |
| Halahakoon, 2024 | 21, 19 | Pramipexole 1mg/day | 0.68 [ 0.04, 1.32] |
| Jocham, 2011 | 16 | Amisulpride 200mg | −0.93 [−1.75, −0.11] |
| Jocham, 2014 | 22 | Amisulpride 400mg | 0.44 [−0.02, 0.90] |
| Kahnt, 2015 | 27, 24 | Amisulpride 400mg | 0.24 [−0.31, 0.79] |
| Murray, 2019 | 18, 18 | Amisulpride 400mg | −0.04 [−0.69, 0.62] |
| Mikus, 2022 | 38, 35 | Amisulpride 400mg | 0.38 [−0.08, 0.84] |
| Cremer, 2023 | 23, 22 | Amisulpride 400mg | 0.59 [−0.00, 1.19] |
| Admon, 2017 | 20, 18 | Amisulpride 50mg | −0.26 [−0.90, 0.38] |
| Frank, 2006 | 19, 20 | Haloperidol 2mg | −0.37 [−1.00, 0.26] |
| Diederen, 2017 | 19, 19 | Sulpiride 600mg | 0.58 [−0.07, 1.23] |
| Haarsma, 2021 | 20, 20 | Sulpiride 600mg | 0.71 [ 0.07, 1.35] |
| Eisenegger, 2014 | 41, 35 | Sulpiride 800mg | −0.05 [−0.50, 0.40] |
| Grob, 2012 | 28, 28 | AMPT 40mg/kg | −0.36 [−0.88, 0.17] |
| Cox, 2015 | 10, 10 | Phe / Tyr depletion 100g | 0.00 [−0.88, 0.88] |
| Random−effects meta−analysis: | | | 0.10 [−0.08, 0.27] |

b

| Study | N(Drug, Pla) | Drug, dose | SMD [95% CI] |
|---|---|---|---|
| Walsh, 2018 | 20, 20 | Bupropion 150mg | 0.37 [−0.25, 1.00] |
| Bernacer, 2013 | 17, 17 | Methamphetamine 0.3mg/kg | −0.01 [−0.68, 0.66] |
| Bellebaum, 2017 | 16, 20 | Modafinil 200mg | −0.52 [−1.19, 0.14] |
| Pessiglione, 2006 | 13, 13 | L−dopa 100mg+25mg | 0.42 [−0.35, 1.20] |
| Halahakoon, 2024 | 21, 19 | Pramipexole 1mg/day | −0.63 [−1.27, 0.00] |
| Jocham, 2011 | 16 | Amisulpride 200mg | −0.08 [−0.92, 0.77] |
| Jocham, 2014 | 22 | Amisulpride 400mg | 0.58 [ 0.07, 1.08] |
| Murray, 2019 | 18, 18 | Amisulpride 400mg | −0.51 [−1.17, 0.16] |
| Admon, 2017 | 20, 18 | Amisulpride 50mg | −0.54 [−1.19, 0.11] |
| Frank, 2006 | 19, 20 | Haloperidol 2mg | 0.38 [−0.25, 1.01] |
| Eisenegger, 2014 | 41, 35 | Sulpiride 800mg | 0.22 [−0.23, 0.67] |
| Cox, 2015 | 10 | Phe / Tyr depletion 100g | −0.91 [−1.95, 0.13] |
| Kelm, 2015 | 16 | Phe / Tyr depletion 68.7g | −0.22 [−0.78, 0.33] |
| Random−effects meta−analysis: | | | −0.07 [−0.32, 0.19] |

c

| Study | N(Drug, Pla) | Drug, dose | SMD [95% CI] |
|---|---|---|---|
| de Wit, 2002 | 36, 36 | D−amphetamine 20mg | −0.21 [−0.67, 0.25] |
| Wardle, 2011 | 17, 17 | D−amphetamine 20mg | −0.64 [−1.33, 0.05] |
| Soder, 2021 | 28, 28 | D−amphetamine 20mg | −0.33 [−0.86, 0.20] |
| Westbrook, 2020 | 46 | Methylphenidate 20mg | −0.19 [−0.48, 0.10] |
| Daood, 2022 | 57 | Methylphenidate 20mg | −0.11 [−0.21, −0.01] |
| Pine, 2010 | 13 | L−dopa 150mg | 0.70 [ 0.06, 1.34] |
| Smith, 2024 | 76, 76 | L−dopa 150mg | −0.14 [−0.46, 0.18] |
| Kayser, 2012 | 23 | Tolcapone 200mg | −0.11 [−0.20, −0.01] |
| Weber, 2016 | 40, 40 | Amisulpride 400mg | 0.58 [ 0.13, 1.02] |
| Wagner, 2020 | 23, 26 | Haloperidol 2mg | 0.53 [−0.04, 1.10] |
| Erfanian Abdoust, 2024 | 62 | Haloperidol 2mg | 0.00 [−0.12, 0.12] |
| Kelm, 2013 | 15 | Phe / Tyr depletion 68.7g | 0.03 [−0.20, 0.26] |
| Random−effects meta−analysis: | | | −0.05 [−0.13, 0.02] |

d

| Study | N(Drug, Pla) | Drug, dose | SMD [95% CI] |
|---|---|---|---|
| Hirschbichler, 2022 | 30 | L−dopa 100mg | 0.51 [ 0.15, 0.87] |
| Rutledge, 2015 | 30 | L−dopa 150mg+37.5mg | 0.31 [ 0.03, 0.60] |
| Rigoli, 2016 | 32 | L−dopa 150mg+37.5mg | 0.12 [−0.08, 0.32] |
| Campbell−Meiklejohn, 2011 | 15, 15 | Pramipexole 0.176μg | −0.31 [−1.03, 0.41] |
| Burke, 2018 | 45, 48 | Amisulpride 400mg | −0.79 [−1.21, −0.37] |
| Random−effects meta−analysis: | | | −0.00 [−0.46, 0.46] |

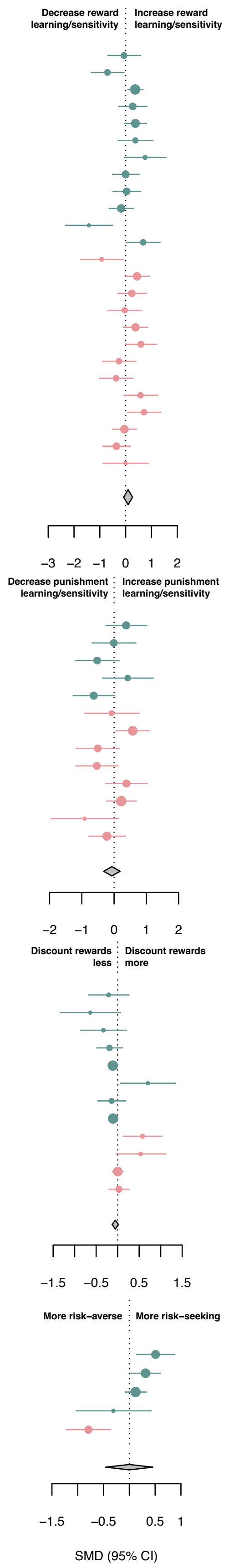


**eFigure 6: The effect of upregulating dopamine on subcomponents according to original drug profile.** The figure shows standardized mean differences (SMDs) between the effect of upregulating dopamine versus placebo on a) reward learning/sensitivity, b) punishment learning/sensitivity, c) reward discounting and d) risk attitude according to the original drug activity profile. Pla indicates placebo. We recoded antagonist effects as if they were agonists (pink SMD point estimates and 95% CI). Green SMD point estimates and 95% CI indicate studies that were coded as original agonists.

a

| Study | N(Drug, Pla) | Drug, dose | SMD [95% CI] |
|---|---|---|---|
| Bernacer, 2013 | 17 | Methamphetamine 0.3mg/kg | −0.70 [−1.33, −0.08] |
| Howlett, 2017 | 19 | Methylphenidate 40mg | 0.27 [−0.26, 0.81] |
| Addicott, 2021 | 22 | Methylphenidate 40mg | 0.38 [−0.03, 0.79] |
| Jansen, 2023 | 30, 30 | L−dopa 100mg+25mg | 0.00 [−0.51, 0.51] |
| Kahnt, 2015 | 27, 24 | Amisulpride 400mg | 0.24 [−0.31, 0.79] |
| Cremer, 2023 | 23, 22 | Amisulpride 400mg | 0.59 [−0.00, 1.19] |
| Diederen, 2017 | 19, 19 | Sulpiride 600mg | 0.58 [−0.07, 1.23] |
| Eisenegger, 2014 | 41, 35 | Sulpiride 800mg | −0.05 [−0.50, 0.40] |
| Random−effects meta−analysis: | | | 0.17 [−0.09, 0.42] |

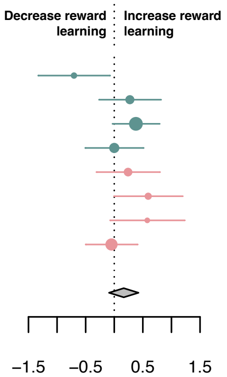


b

| Study | N(Drug, Pla) | Drug, dose | SMD [95% CI] |
|---|---|---|---|
| Bernacer, 2013 | 17, 17 | Methamphetamine 0.3mg/kg | 0.23 [−0.44, 0.90] |
| Addicott, 2021 | 22 | Methylphenidate 40mg | −0.18 [−0.62, 0.27] |
| Mikus, 2022 | 38, 35 | Amisulpride 400mg | 0.38 [−0.08, 0.84] |
| Cremer, 2023 | 23, 22 | Amisulpride 400mg | −0.48 [−1.07, 0.12] |
| Eisenegger, 2014 | 41, 35 | Sulpiride 800mg | 0.68 [ 0.22, 1.14] |
| Random−effects meta−analysis: | | | 0.14 [−0.27, 0.55] |

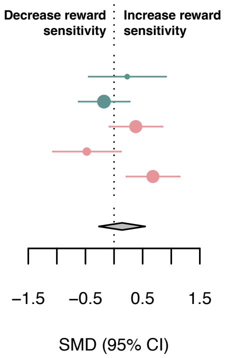


c

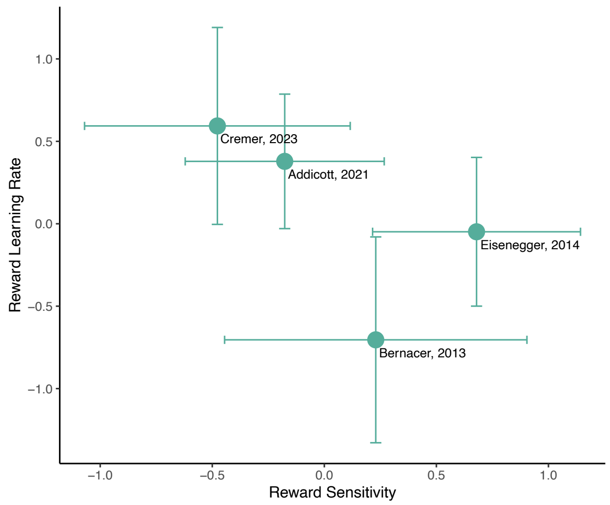


d

| Study | N(Drug, Pla) | Drug, dose | SMD [95% CI] |
|---|---|---|---|
| Bernacer, 2013 | 17, 17 | Methamphetamine 0.3mg/kg | 0.12 [−0.55, 0.79] |
| Eisenegger, 2014 | 41, 35 | Sulpiride 800mg | −0.01 [−0.46, 0.44] |
| Random−effects meta−analysis: | | | 0.03 [−0.35, 0.40] |

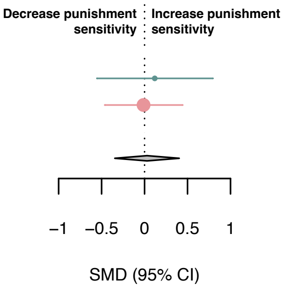


**eFigure 7: The effect of upregulating dopamine on learning rate and sensitivity parameters.** Standardized mean differences (SMDs) between the effect of upregulating dopamine versus placebo on a) reward learning rates, b) reward sensitivity, c) scatter plot of reward learning rate and reward sensitivity SMDs for studies that reported both - error bars represent 95% CI, and d) punishment sensitivity. Pla indicates placebo. We recoded antagonist effects as if they were agonists (pink SMD point estimates and 95% CI). Green SMD point estimates and 95% CI indicate studies that were coded as original agonists.

#### 2.3 Dopamine heterogeneity

There was low-to-medium heterogeneity in overall reward ($\tau^2$=0.04, $I^2$=58.50%) and overall punishment processes ($\tau^2$=0.07, $I^2$=41.74%). Excluding four identified outlier studies[2,27,47,65] reduced heterogeneity to low in overall reward processes ($\tau^2$=0.01; $I^2$=31.39%), but did not substantially affect the pooled effect size (SMD=0.19, 95% CI [0.11, 0.26]).

There was low heterogeneity in the reward learning/sensitivity subcomponent ($\tau^2$=0.05, $I^2$=39.31%; removing two identified outliers[2,27] reduced this heterogeneity further $\tau^2$=0.01, $I^2$=12.94%, which did not significantly affect the pooled effect size (SMD=0.26, 95% CI [0.14, 0.38]). There was moderate-to-substantial heterogeneity in the punishment learning/sensitivity ($\tau^2$=0.11, $I^2$=50.74%), appetitive Pavlovian ($\tau^2$=0.14, $I^2$=67.66%), risk attitude ($\tau^2$=0.22, $I^2$=88.22%), and model-based learning/flexibility subcomponent ($\tau^2$=0.31, $I^2$=84.76%). Removing an identified outlier in the risk attitude subcomponent[66] reduced heterogeneity to $\tau^2$=0.01, $I^2$=32.98%, which resulted in a significant SMD estimate (SMD=0.27, 95% CI [0.08, 0.47]; eFigure 8). There was no-to-low heterogeneity in the reward response vigor ($\tau^2$=0.02, $I^2$=19.18%), reward discounting ($\tau^2$=0.0009, $I^2$=7.35%), and aversive Pavlovian ($\tau^2$<0, $I^2$<0%) subcomponent. No other outliers were detected in either the subcategories or overall sample.

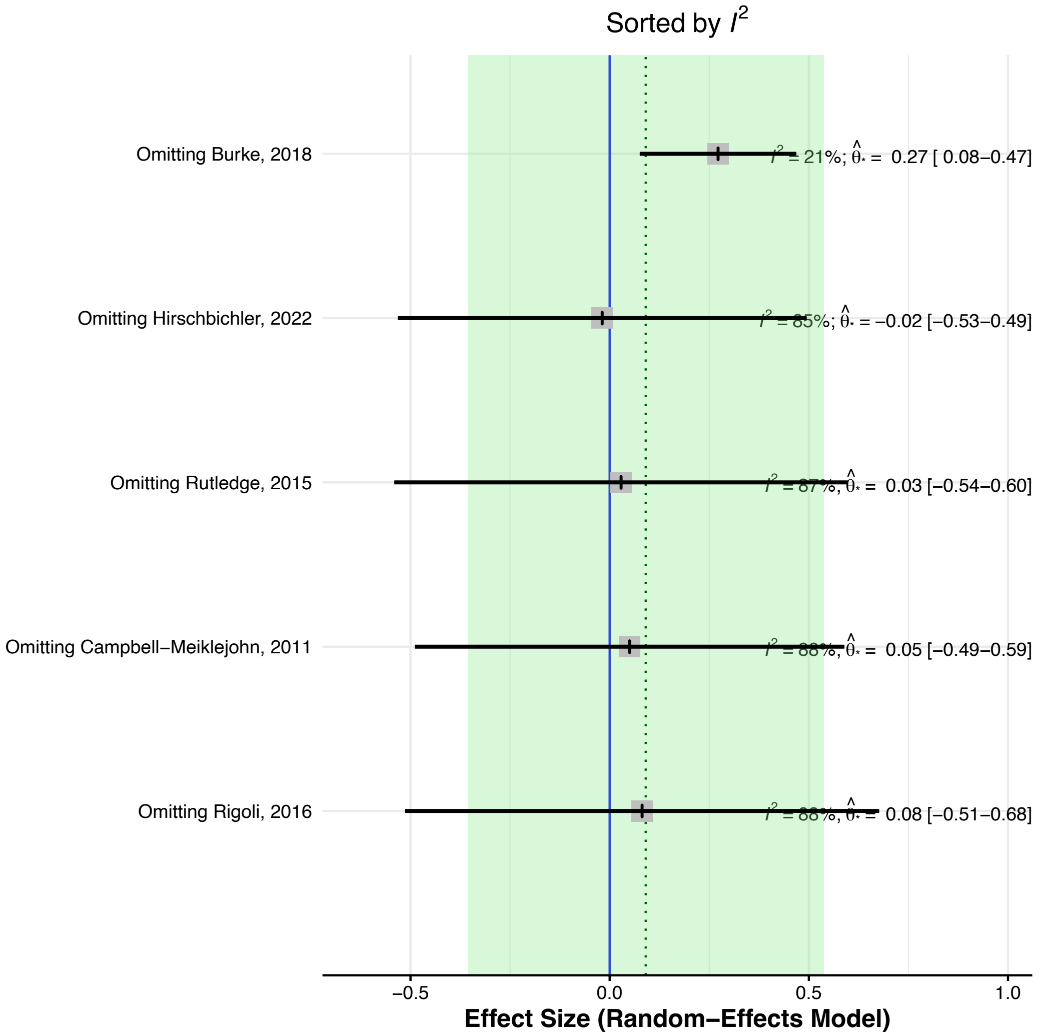


**eFigure 8: Leave-one-out meta-analysis for the effect of dopamine on risk attitude.** It is ordered by heterogeneity (low to high), displaying the recalculated pooled effect size and 95% confidence interval with one study omitted each time. The green shaded area represents the 95% confidence interval of the original pooled effect size, and the dotted line represents the original estimated pooled effect size.

#### 2.4 Additional serotonin results

a

| Study | N(Drug, Pla) | Drug, dose | SMD [95% CI] |
|---|---|---|---|
| Colwell, 2024 | 26, 27 | Fenfluramine 15mg b.i.d. | 0.03 [−0.51, 0.57] |
| Michely, 2022 | 33, 33 | Citalopram 20mg/day | −0.66 [−1.16, −0.17] |
| Guitart−Masip, 2012 | 16, 20 | Citalopram 24mg oral | −0.26 [−0.92, 0.40] |
| Beierholm, 2013 | 30, 30 | Citalopram 24mg oral | 0.26 [−0.25, 0.76] |
| Jepma, 2010 | 16, 17 | Citalopram 30mg | −0.04 [−0.72, 0.64] |
| Guitart−Masip, 2014 | 27, 29 | Citalopram 30mg | 0.28 [−0.24, 0.81] |
| Luo, 2024 | 32, 33 | Escitalopram 20mg | −3.28 [−4.02, −2.54] |
| Crean, 2002 | 20, 20 | TD | 0.29 [−0.33, 0.91] |
| Cools, 2005 | 10, 12 | TD | 1.20 [ 0.29, 2.12] |
| Roiser, 2006 | 30 | TD | 0.16 [−0.29, 0.61] |
| Finger, 2007 | 16, 19 | TD | 0.59 [−0.09, 1.27] |
| Tanaka, 2007 | 12, 12 | TD | −0.02 [−0.82, 0.78] |
| Cools, 2008 | 12 | TD | −0.02 [−0.49, 0.46] |
| Schweighofer, 2008 | 20 | TD | 0.77 [ 0.13, 1.41] |
| Robinson, 2012 | 21, 20 | TD | −0.31 [−0.93, 0.30] |
| Seymour, 2012 | 15, 15 | TD | 1.02 [ 0.26, 1.78] |
| Geurts, 2013 | 45 | TD | 0.02 [−0.32, 0.35] |
| Worbe, 2014 | 22, 22 | TD | 0.24 [−0.36, 0.83] |
| Hebart, 2015 | 34, 34 | TD | −0.00 [−0.48, 0.48] |
| Steding, 2023 | 24 | TD | 0.08 [−0.12, 0.27] |
| Gilger, 2024 | 98, 98 | TD | 0.03 [−0.25, 0.31] |
| Random−effects meta−analysis: | | | 0.02 [−0.33, 0.36] |

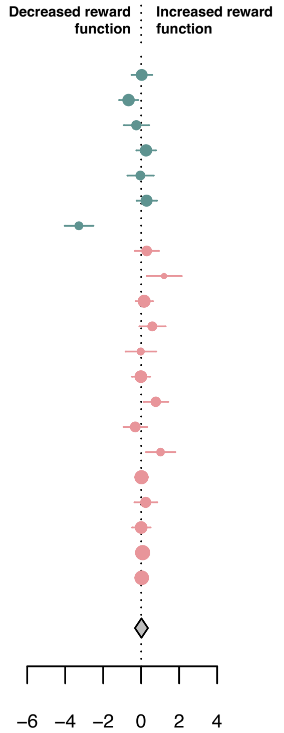


b

| Study | N(Drug, Pla) | Drug, dose | SMD [95% CI] |
|---|---|---|---|
| Colwell, 2024 | 26, 27 | Fenfluramine 15mg b.i.d. | 0.41 [−0.13, 0.95] |
| Michely, 2022 | 33, 33 | Citalopram 20mg/day | 0.49 [ 0.00, 0.98] |
| Guitart−Masip, 2012 | 16, 20 | Citalopram 24mg oral | −0.00 [−0.66, 0.66] |
| Guitart−Masip, 2014 | 27, 29 | Citalopram 30mg | −0.41 [−0.94, 0.12] |
| Luo, 2024 | 32, 33 | Escitalopram 20mg | 0.33 [−0.16, 0.82] |
| Cools, 2008 | 12 | TD | 0.59 [ 0.04, 1.14] |
| Crockett, 2012 | 24 | TD | 0.65 [ 0.03, 1.27] |
| Robinson, 2012 | 21, 20 | TD | 0.65 [ 0.02, 1.28] |
| Seymour, 2012 | 15, 15 | TD | −0.20 [−0.92, 0.51] |
| Geurts, 2013 | 45 | TD | 0.48 [ 0.05, 0.91] |
| Hebart, 2015 | 34, 34 | TD | −0.57 [−1.06, −0.09] |
| Random−effects meta−analysis: | | | 0.22 [−0.04, 0.49] |

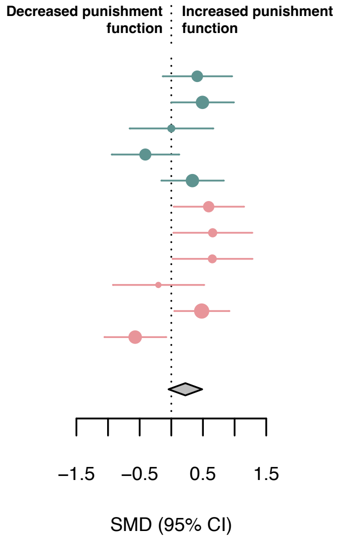


**eFigure 9: The effect of upregulating serotonin on overall reward and punishment.** Standardized mean differences (SMDs) between the effect of upregulating serotonin versus placebo on a) overall reward function, and b) overall punishment function. Pla indicates placebo. We recoded antagonist effects as if they were agonists (pink SMD point estimates and 95% CI). Green SMD point estimates and 95% CI indicate studies that were coded as original agonists. TD: tryptophan depletion.

a

| Study | N(Drug, Pla) | Drug, dose | SMD [95% CI] |
|---|---|---|---|
| Macoveanu, 2014 | 13, 16 | Fluoxetine 20/40mg/day | 0.38 [−0.36, 1.12] |
| Murphy, 2009 | 15, 15 | L−tryptophan 3g/day | −0.50 [−1.23, 0.22] |
| Anderson, 2003 | 15, 13 | TD | 0.03 [−0.71, 0.77] |
| Rogers, 2003 | 18, 18 | TD | −0.21 [−0.86, 0.45] |
| Campbell−Meiklejohn, 2011 | 17, 17 | TD | 1.00 [ 0.28, 1.71] |
| Faulkner, 2017 | 26 | TD | 0.37 [−0.04, 0.78] |
| Random−effects meta−analysis: | | | 0.19 [−0.21, 0.59] |

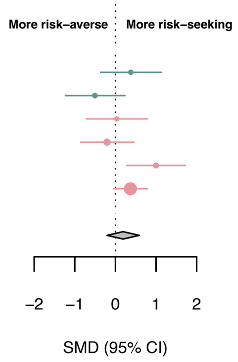


b

| Study | N(Drug, Pla) | Drug, dose | SMD [95% CI] |
|---|---|---|---|
| Chamberlain, 2006 | 20, 20 | Citalopram 30mg | −0.18 [−0.80, 0.44] |
| Langley, 2023 | 32, 34 | Escitalopram 20mg /day | −0.26 [−0.75, 0.22] |
| Skandali, 2018 | 31, 33 | Escitalopram 20mg | −0.28 [−0.77, 0.22] |
| Thirkettle, 2019 | 22, 22 | Tryptophan 0.8g | 0.01 [−0.58, 0.60] |
| Evers, 2004 | 15, 15 | TD | 0.29 [−0.43, 1.01] |
| Cools, 2008 | 12 | TD | 0.09 [−0.34, 0.51] |
| Worbe, 2015 | 18, 18 | TD | 0.78 [ 0.10, 1.46] |
| Worbe, 2016 | 22, 22 | TD | 0.79 [ 0.18, 1.41] |
| Kanen, 2020 | 30, 32 | TD | −0.04 [−0.54, 0.46] |
| Gilger, 2024 | 98, 98 | TD | −0.02 [−0.30, 0.26] |
| Random−effects meta−analysis: | | | 0.06 [−0.13, 0.26] |

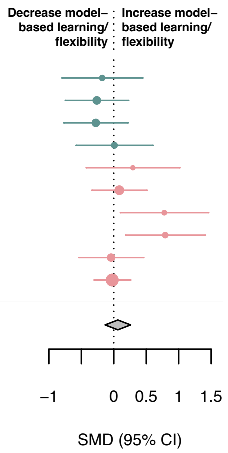


**eFigure 10: The effect of upregulating serotonin on risk attitude and model-based learning/flexibility.** Standardized mean differences (SMDs) between the effect of upregulating serotonin versus placebo on a) risk attitude, and b) model-based learning/flexibility. Pla indicates placebo. We recoded antagonist effects as if they were agonists (pink SMD point estimates and 95% CI). Green SMD point estimates and 95% CI indicate studies that were coded as original agonists. TD: tryptophan depletion.

| Study | N(Drug, Pla) | Drug, dose | SMD [95% CI] |
|---|---|---|---|
| Cools, 2008 | 12 | TD | 0.85 [ 0.19, 1.51] |
| Crockett, 2009 | 22 | TD | 0.49 [ 0.04, 0.93] |
| Crockett, 2012 | 24 | TD | 0.48 [ 0.06, 0.90] |
| Geurts, 2013 | 45 | TD | 0.35 [ 0.05, 0.65] |
| Gaber, 2015 | 24 | TD | 0.11 [−0.29, 0.52] |
| Helmbold, 2015 | 18 | TD | 0.22 [−0.24, 0.69] |
| Random−effects meta−analysis: | | | 0.36 [ 0.20, 0.53] |

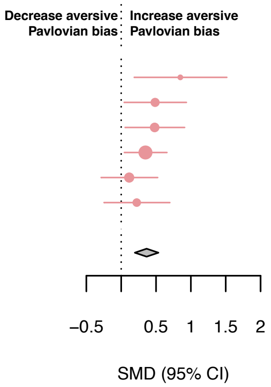


**eFigure 11: The effect of upregulating serotonin on aversive Pavlovian biases in within-subject studies.** Standardized mean differences (SMDs) between the effect of upregulating serotonin versus placebo on aversive Pavlovian bias, only including within-subject studies. These SMDs reflect Cohen's $d_z$. Pla indicates placebo. We recoded antagonist effects as if they were agonists (pink SMD point estimates and 95% CI). TD: tryptophan depletion.

a

| Study | N(Drug, Pla) | Drug, dose | SMD [95% CI] |
|---|---|---|---|
| Colwell, 2024 | 26, 27 | Fenfluramine 15mg b.i.d. | 0.03 [−0.51, 0.57] |
| Michely, 2022 | 33, 33 | Citalopram 20mg/day | −0.66 [−1.16, −0.17] |
| Luo, 2024 | 32, 33 | Escitalopram 20mg | −3.28 [−4.02, −2.54] |
| Random−effects meta−analysis: | | | −1.29 [−3.25, 0.67] |

b

| Study | N(Drug, Pla) | Drug, dose | SMD [95% CI] |
|---|---|---|---|
| Colwell, 2024 | 26, 27 | Fenfluramine 15mg b.i.d. | −0.06 [−0.60, 0.48] |
| Jepma, 2010 | 16, 17 | Citalopram 30mg | −0.04 [−0.72, 0.64] |
| Seymour, 2012 | 15, 15 | TD | 1.02 [ 0.26, 1.78] |
| Random−effects meta−analysis: | | | 0.27 [−0.40, 0.94] |

c

| Study | N(Drug, Pla) | Drug, dose | SMD [95% CI] |
|---|---|---|---|
| Colwell, 2024 | 26, 27 | Fenfluramine 15mg b.i.d. | 0.41 [−0.13, 0.95] |
| Michely, 2022 | 33, 33 | Citalopram 20mg/day | 0.49 [ 0.00, 0.98] |
| Luo, 2024 | 32, 33 | Escitalopram 20mg | 0.33 [−0.16, 0.82] |
| Random−effects meta−analysis: | | | 0.41 [ 0.12, 0.70] |

d

| Study | N(Drug, Pla) | Drug, dose | SMD [95% CI] |
|---|---|---|---|
| Colwell, 2024 | 26, 27 | Fenfluramine 15mg b.i.d. | −0.57 [−1.12, −0.02] |
| Seymour, 2012 | 15, 15 | TD | −0.20 [−0.92, 0.51] |
| Random−effects meta−analysis: | | | −0.44 [−0.87, −0.00] |

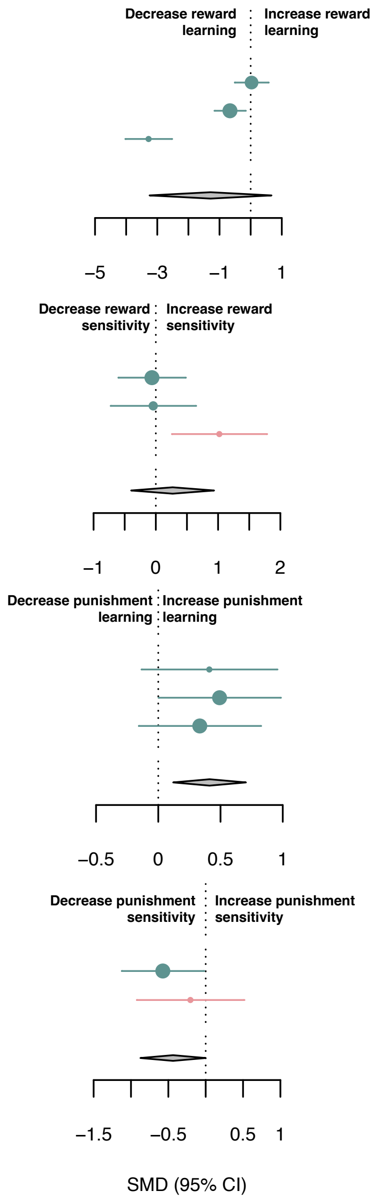


**eFigure 12: The effect of upregulating serotonin on learning rate and sensitivity parameters.** Standardized mean differences (SMDs) between the effect of upregulating serotonin versus placebo on a) reward learning rates, b) reward sensitivity, c) punishment learning rates, and d) punishment sensitivity. Pla indicates placebo. We recoded antagonist effects as if they were agonists (pink SMD point estimates and 95% CI). Green SMD point estimates and 95% CI indicate studies that were coded as original agonists. TD: tryptophan depletion.

#### 2.5 Serotonin heterogeneity

Between-subject heterogeneity in overall reward ($\tau^2$= 0.55, $I^2$= 90.44%) and punishment ($\tau^2$= 0.12, $I^2$= 61.32%) processes were moderate-to-substantial. Removing one outlier study[67] in the overall reward category reduced heterogeneity to low, explaining a substantial part of the heterogeneity ($\tau^2$= 0.02, $I^2$= 28.06%), resulting in a small positive but nonsignificant pooled effect size (SMD=0.11; 95% CI, [-0.02, 0.24]). Removing one outlier[18] in overall punishment function reduced the heterogeneity to low ($\tau^2$ = 0.05, $I^2$= 37.02%), which increased the overall effect size and made it significant (SMD=0.32; 95% CI, [0.10, 0.54]; eFigure 13).

There was no heterogeneity in the punishment learning/sensitivity ($\tau^2$<0, $I^2$= 0.01%), appetitive Pavlovian ($\tau^2$<0, $I^2$<0%), reward response vigor ($\tau^2$=0.01, $I^2$=17.37%) and reward discounting ($\tau^2$<0, $I^2$<0%) subcomponent. There was low-to-moderate heterogeneity in the model-based learning/flexibility ($\tau^2$=0.032, $I^2$=33.40%) and risk attitude subcomponent ($\tau^2$= 0.13, $I^2$=55.67%) and substantial heterogeneity in the reward learning/sensitivity ($\tau^2$=1.30, $I^2$=94.70%) and aversive Pavlovian category ($\tau^2$=0.21, $I^2$=72.66%). Removing one identified outlier[67] in the reward learning/sensitivity subcomponent reduced the heterogeneity somewhat ($\tau^2$=0.13, $I^2$=64.95%) but did not affect the pooled effect size (SMD=0.03; 95% CI [-0.28, 0.35]). No other outliers were detected.

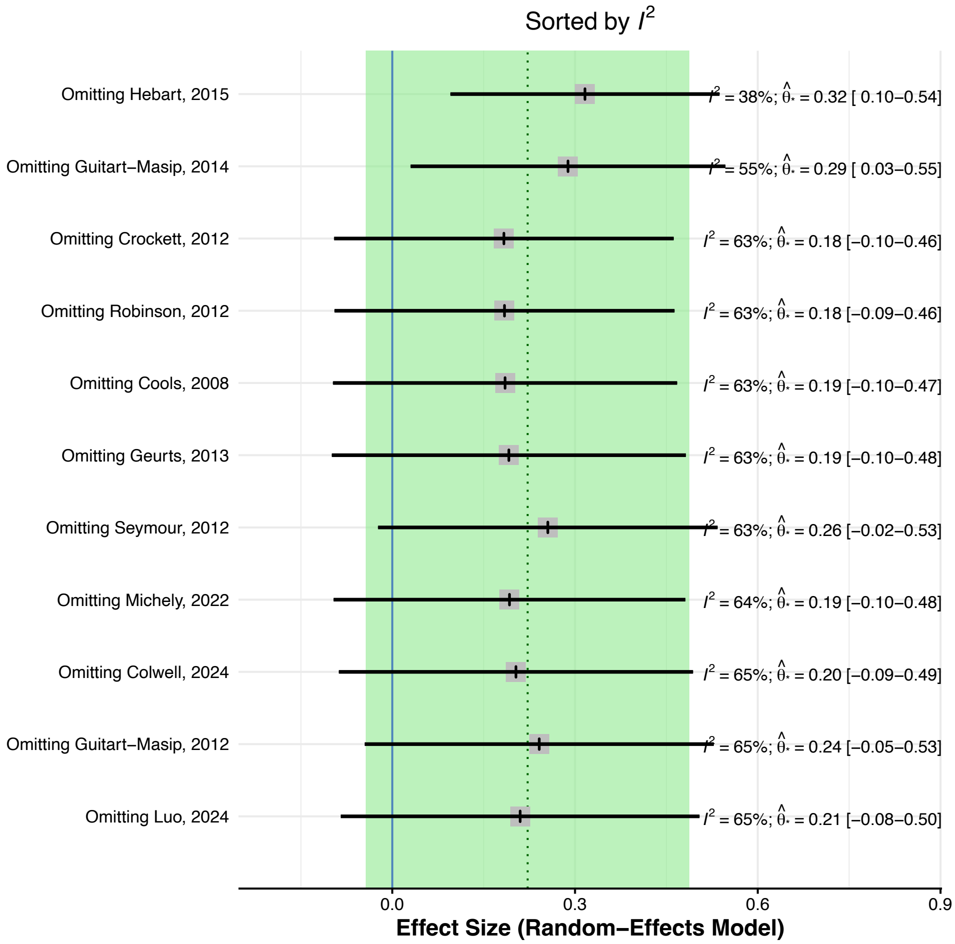


**eFigure 13: Leave-one-out meta-analysis for the effect of serotonin on overall punishment function.** It is ordered by heterogeneity (low to high), displaying the recalculated pooled effect size and 95% confidence interval with one study omitted each time. The green shaded area represents the 95% confidence interval of the original pooled effect size, and the dotted line represents the original estimated pooled effect size.

#### 2.6 Moderator analyses

Several moderators were used to assess potential sources of bias in overall reward and punishment processing and subcomponents with sufficient number of studies (N>9) and substantial heterogeneity. These included reward and punishment learning/sensitivity and reward discounting components in the dopamine domain and the model-based learning/flexibility subcomponent in the serotonin domain. Task type was only assessed for the subcomponents since these would likely just recapitulate the subcomponents in the overall reward/punishment domains.

##### 2.6.1 Dopamine

eTable 3 lists the number of studies included for all the DA moderator analyses. None of the moderators were significant predictors in the overall reward/punishment categories (proportion of female participants: reward p=0.98; punishment p=0.998; drug regimen - single vs. multiple drug administrations: reward p=0.40, not assessed in overall punishment due to too few studies with multiple drug administrations; drug type - agonist vs. antagonist: reward p=0.73; punishment p=0.75; more fine-grained drug activity classifications: reward p=0.64; punishment p=0.12; overall study age: reward p=0.91; punishment p=0.82).

Drug type was a significant predictor of dopamine in the reward discounting subcomponent ($R^2$= 13.88%, $I^2$= 6.45%, p=0.03). Studies that originally used an agonist (10 studies) had a small significant decreasing impact on reward discounting (SMD=-0.10, 95% CI [-0.16, -0.03]) compared with studies that originally used an antagonist (2 studies; note they are recoded to represent agonist), which had a small, non-significant, increasing effect on reward discounting (SMD=0.14, 95% CI [-0.06, 0.35]). It is not immediately clear what might be driving the difference between agonist and antagonist here. It is possible that this might stem from interactions with baseline dopamine levels[68] or task differences. All the antagonist studies used temporal delay discounting tasks, which we in a below exploratory analysis find are less impacted by dopamine here. This might also align with a recent theory suggesting that dopamine, particularly antagonists, might have differential effects on temporal, compared with, effort delay discounting[69] processes. However, due to the exploratory nature of this analysis, future research is needed to determine if different drug activity types do indeed have different reward discounting effects, particularly given the limited number of antagonist studies included here.

Drug type did not moderate reward or punishment learning/sensitivity (reward: p=0.60; punishment: p=0.92). More fine-grained drug type (reward: p=0.48; punishment: p=0.24; reward discounting: p=0.40), age (reward: p=0.41; too few punishment and reward discounting studies to assess) or proportion of female participants (reward: p=0.76; punishment: p=0.997; reward discounting: p=0.17) did not moderate the reward/punishment learning/sensitivity or reward discounting results. There was only one study in the reward and punishment learning/sensitivity subcomponents that used multiple drug administrations and none in the reward discounting subcomponent. This moderator was therefore not evaluated here.

Task type did not moderate reward (p=0.26) or punishment learning/sensitivity (p=0.41). As an additional exploratory analysis, we examined if the results in the reward discounting category might differ depending on the task type. This was explored based on previous reviews suggesting that dopamine might differentially affect effort and temporal discounting[69]. Although effort versus temporal delay discounting studies were not significantly different from each other (p=0.09), the effect of dopamine on reward discounting was only significant in the effort discounting paradigms (SMD=-0.28, 95% CI [-0.52, -0.03]; 3 studies) but not in the temporal delay discounting tasks (SMD=-0.06, 95% CI [-0.13, 0.02]; 9 studies). However, due to the lack of significant difference of dopamine between these tasks, we do not interpret this further but suggest this as an area for future research to resolve.

| Moderator | Overall reward | Overall punishment | Reward learning/sensitivity | Punishment learning/sensitivity | Reward discounting |
|---|---|---|---|---|---|
| Drug type | DA+=32<br>DA-=17 | DA+=11<br>DA-=6 | DA+=14<br>DA-=11 | DA+=8<br>DA-=5 | DA+=10<br>DA-=2 |
| Specific drug type | D2-=16<br>DAP=11<br>DARI=11,<br>other=11 | D2-=6<br>other=11 | D2-=11<br>DAP=4<br>DARI=6<br>other=4 | D2-=6<br>other=7 | D2-=3<br>DAP=2<br>DARI=5<br>other=2 |
| Task type | - | - | PILT=10<br>PST=5<br>Other=10 | PILT/Other=8<br>PST=5 | DD=9<br>Effort=3 |
| Drug regimen | multiple=3<br>single=46 | multiple=1<br>single=16 | multiple=1<br>single=24 | multiple=1<br>single=12 | single=12 |
| Age | N=34 | N=10 | N=16 | N=8 | N=8 |
| % Female | N=41 | N=13 | N=20 | N=11 | N=10 |

**eTable 3: Number of studies for each moderator in the dopamine (DA) domain.** DA+: agonist, DA-: antagonist, D2-: D2-receptor antagonist, DAP: DA precursor, DARI: DA reuptake inhibitor, PILT: Probabilistic instrumental learning task, PST: Probabilistic selection task, DD: delay discounting task, Effort: effort discounting task

##### 2.6.2 Serotonin

eTable 4 describes the number of studies included for all the serotonin moderator analyses. A subgroup analysis indicated that drug type significantly moderated the effect sizes in overall reward processing ($R^2$=17.82%, $I^2$=88.49%, p=0.03). However, further analysis showed that the individual effect sizes for both the antagonist (SMD=0.26, 95% CI [-0.12, 0.65]) and agonist (SMD=-0.48, 95% CI [-1.03, 0.06]) subgroups were not significant, indicating that the overall significant moderation effect is driven by differences in directionality rather than by significant effects within each group. This effect is likely driven by the previously identified outlier[67]. Removing this outlier, drug activity is no longer a significant predictor (p=0.13).

The same was true for drug type moderating effect sizes in the model-based/flexibility subcategory ($R^2$=39.04%, $I^2$=22.92%, p=0.046), with neither the agonist (SMD=-0.19, 95% CI [-0.49, 0.11]) or antagonist (SMD=0.19, 95% CI [-0.03, 0.42]) subgroup having a significant overall effect on model-based learning. Drug type was not a significant moderator of the effect of serotonin on overall punishment function (p=0.76). Subgroup analyses with more fine-grained drug activity was not conducted since these included virtually the same groupings as the agonist/antagonist subgroup analysis. Drug regimen (single vs multiple drug administrations) did not significantly moderate the overall reward (p=0.53) or punishment (p=0.42) processes. There was only one study with multiple drug administrations in the model-based/flexibility subcomponent and thus this moderator was not assessed here. Neither age nor proportion of female participants were significant predictors in the overall reward category (age: p=0.80, %female participants: p=0.33) or in the model-based/flexibility subcomponent (too few studies to examine age; %female participants: p=0.27). There were too few studies to examine age and %female participants in the overall punishment category. Task type was not a significant moderator of model-based learning/flexibility (p=0.11).

| Moderator | Overall reward | Overall punishment | Model-based learning/flexibility |
|---|---|---|---|
| Drug type | 5HT+=7<br>5HT-=14 | 5HT+=5<br>5HT-=6 | 5HT+=4<br>5HT-=6 |
| Task type | - | - | two-step=3<br>reversal=6<br>other=1 |
| Drug regimen | multiple=2<br>single=19 | multiple=2<br>single=9 | multiple=1<br>single=9 |
| Age | N=15 | N=7 | N=9 |
| % Female | N=17 | N=6 | N=10 |

**eTable 4: Number of studies for each moderator in the serotonin (5HT) domain.** 5HT+: agonist, 5HT-: antagonist.

#### 2.7 Publication bias

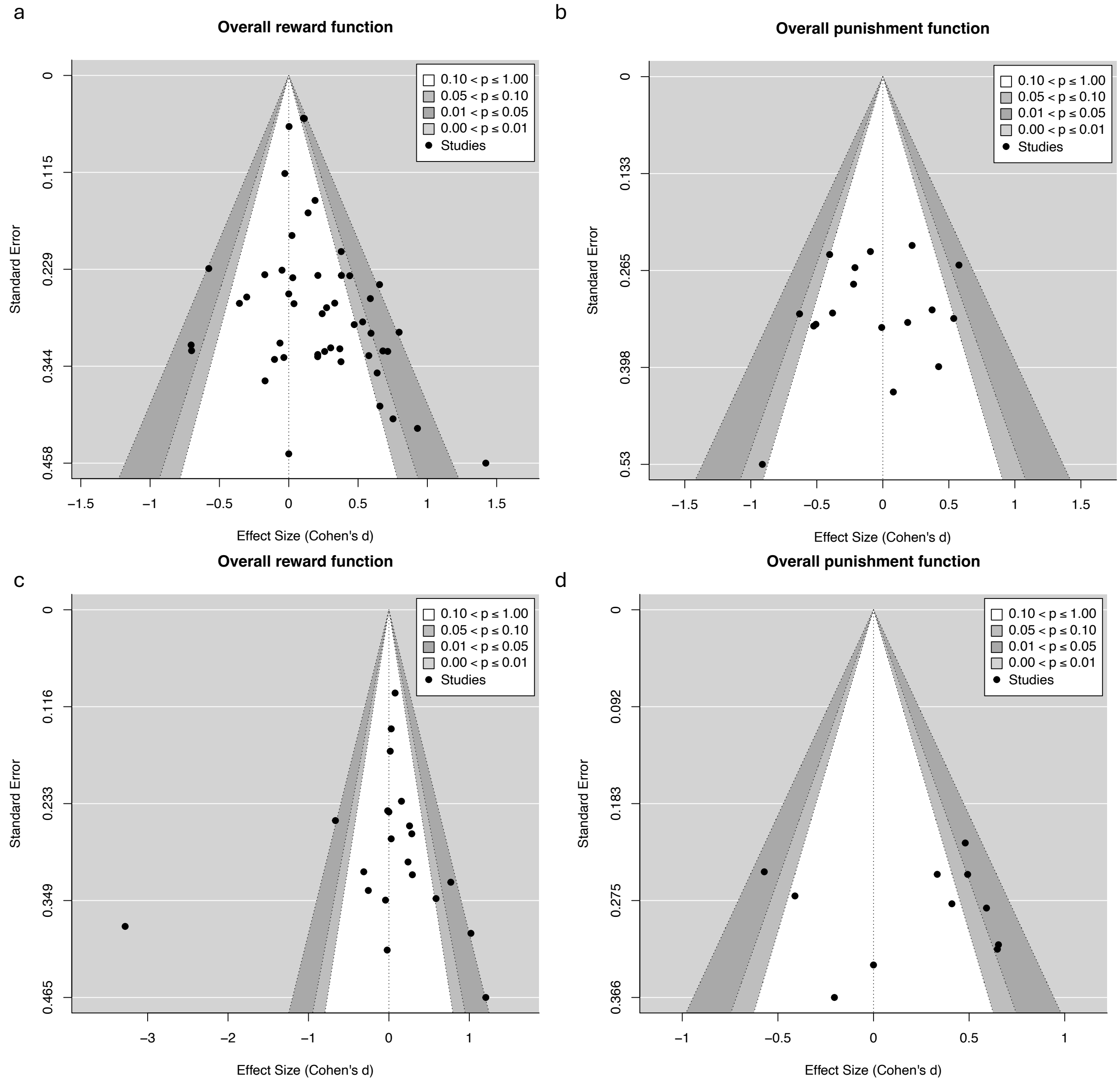


**eFigure 14: Funnel plots of overall reward and dopamine function.** Funnel plots of dopamine on a) overall reward function, b) overall punishment function, and funnel plots of serotonin on c) overall reward function, and d) overall punishment function.

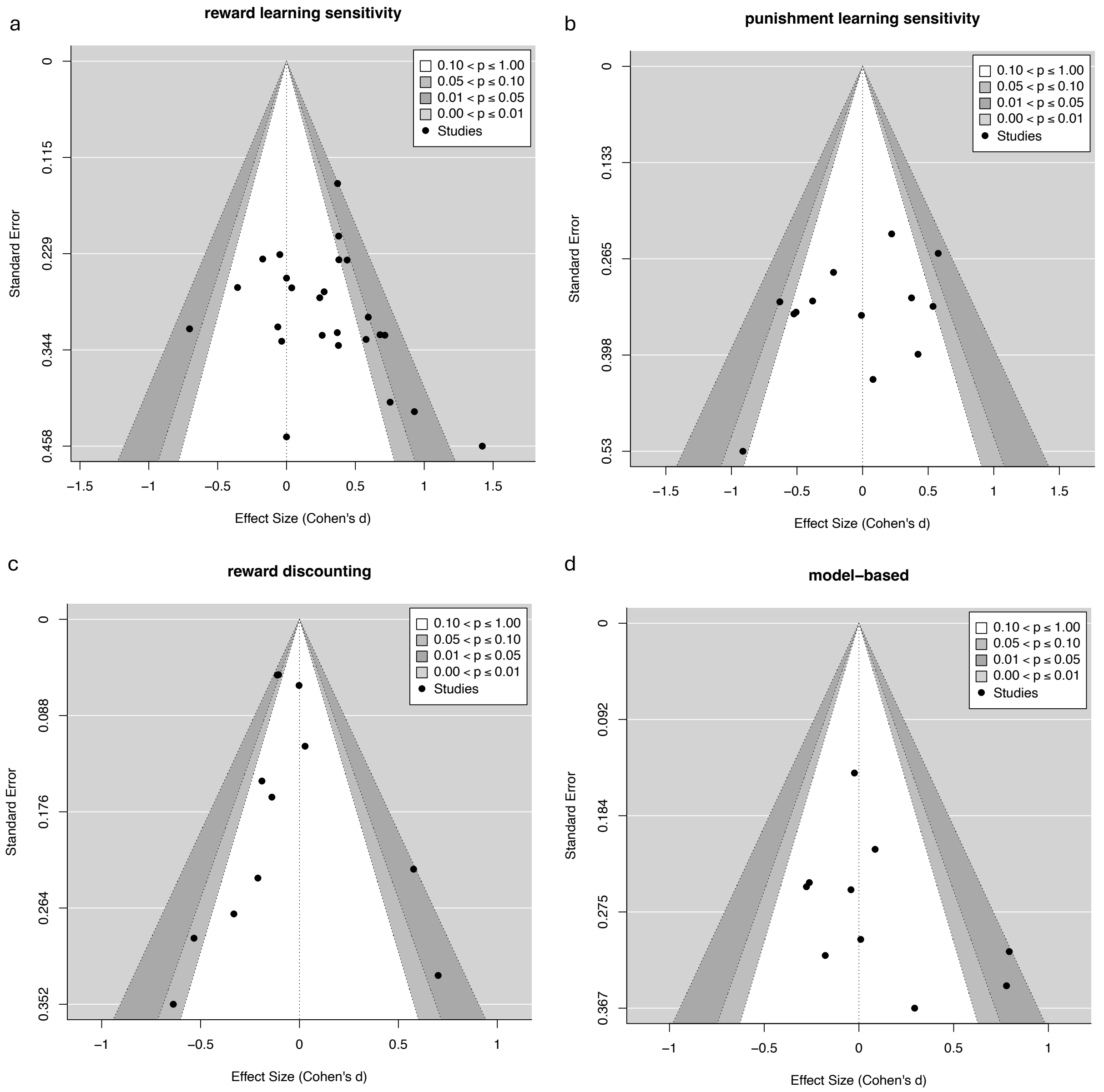


**eFigure 15: Funnel plots of subcomponents.** Funnel plots of dopamine on a) reward learning/sensitivity, b) punishment learning/sensitivity, c) reward discounting, and funnel plots of serotonin on d) model-based learning/flexibility.

#### 2.8 Additional supplementary analyses

We note that olanzapine has a more complex drug activity profile than the other drug compounds included in the dopamine reward response vigor subcategory. Given the broader receptor activity profile of olanzapine, we performed a sensitivity analysis excluding this study to ensure it did not unduly influence the vigor results. The exclusion of this study[45] did not substantially alter the overall findings or conclusions (SMD = 0.30, 95% CI [0.07, 0.52]).

We categorized discounting tasks in line with previous meta-analyses where they are referred to as option valuation and categorized within the overall reward function[70,71]. This decision was further based on the fact that reward discounting paradigms typically involve rewards represented as points or money, aligning with other reward-processing tasks in this category. In contrast, tasks in the overall punishment function category primarily involve explicit losses of points or money, whereas discounting tasks incorporate aversive elements such as time delays or effort costs. Because reward discounting is typically interpreted in terms of how much a reward is valued rather than as a direct punishment process, we believe that this categorization provides a clearer theoretical interpretation. However, we recognize that the classification of reward discounting tasks is not entirely straightforward and could also be considered part of an aversive process. Grouping reward discounting as a punishment processing task, resulted in a small, non-meaningful, negative dopamine effect on overall punishment function (SMD=-0.07, 95%CI [-0.17, 0.03]), and a non-meaningful positive serotonin effect on overall punishment function (SMD=0.09, 95%CI [-0.16, 0.33]).

| Study | Category | Drug, Dose | N drug | N placebo | Original design | Final design | Task | Outcome | Study Age | Study Female Proportion | Drug regimen |
|---|---|---|---|---|---|---|---|---|---|---|---|
| Addicott, 2021[72] | R L/S | methylphenidate 40mg | 22 | 22 | w | w | restless bandit | learning rate | 27.7 | 0.57 | 1 |
| Admon, 2017*[73] | R L/S | amisulpride 50mg | 20 | 18 | bw | bw | PST | choose A | 25.9 | 0.77 | 1 |
| Bellebaum, 2017[1] | R L/S | modafinil 200mg | 16 | 20 | bw | bw | PST | choose A | 24.85 | 0 | 1 |
| Bernacer, 2013[2] | R L/S | methamphetamine 0.3mg/kg | 17 | 17 | w | w | PILT | learning rate | 25.3 | 0.41 | 1 |
| Chakroun, 2023[74] | R L/S | L-dopa 150mg+37.5mg | 31 | 31 | w | w | PILT | accuracy | 26.85 | 0 | 1 |
| Cox, 2015[7] | R L/S | Phe / Tyr depletion 100g | 10 | 10 | w† | bw | PST | choose A/C | NA | NA | 1 |
| Cremer, 2023[75] | R L/S | amisulpride 400mg | 23 | 22 | bw | bw | foraging | learning rate | NA | 0.47 | 1 |
| Diederen, 2017[76] | R L/S | sulpiride 600mg | 19 | 19 | bw | bw | observational reversal | learning rate | 24.4 | 0.47 | 1 |
| Eisenegger, 2014[13] | R L/S | sulpiride 800mg | 41 | 35 | bw | bw | PILT | learning rate | NA | 0 | 1 |
| Frank, 2006*[14] | R L/S | haloperidol 2mg | 19 | 20 | w† | bw | PST | choose A | 21 | 0.54 | 1 |
| Grob, 2012[77] | R L/S | AMPT 40mg/kg | 28 | 28 | w† | bw | PRT | response bias block 3 | 25.8 | 1 | 1 |
| Haarsma, 2021[78] | R L/S | sulpiride 600mg | 20 | 20 | bw | bw | reward prediction | accuracy | 24.35 | 0.42 | 1 |
| Halahakoon, 2024[79] | R L/S | pramipexole 1mg/day | 21 | 19 | bw | bw | PILT | accuracy change pre-post intervention | 23.5 | 0.5 | >1 |
| Howlett, 2017[80] | R L/S | methylphenidate 40mg | 19 | 19 | w | w | random dot based PILT | learning rate | NA | 0 | 1 |
| Jansen, 2023[81] | R L/S | L-dopa 100mg+25mg | 30 | 30 | w† | bw | PILT | learning rate | 22.8 | 0 | 1 |
| Jocham, 2011*[53] | R L/S | amisulpride 200mg | 16 | 16 | w | w | PST | p(correct) win-win | 26.13 | 0 | 1 |
| Jocham, 2014[19] | R L/S | amisulpride 400mg | 22 | 22 | w | w | PILT | final accuracy | NA | 0 | 1 |
| Kahnt, 2015[82] | R L/S | amisulpride 400mg | 27 | 24 | bw | bw | noninstrumental outcome prediction | learning rate | 22.4 | 0 | 1 |
| Mikus, 2022[24] | R L/S | amisulpride 400mg | 38 | 35 | bw | bw | Two-step task | inverse temperature | NA | NA | 1 |
| Murray, 2019[83] | R L/S | amisulpride 400mg | 18 | 18 | w† | bw | PILT | number correct choice | 32.1 | 0.17 | 1 |
| Pessiglione, 2006[26] | R L/S | L-dopa 100mg+25mg | 13 | 13 | bw | bw | PILT | accuracy | NA | NA | 1 |
| Pizzagalli, 2008*[27] | R L/S | pramipexole 0.5mg | 11 | 13 | bw | bw | PRT | response bias block 3-1 | NA | NA | 1 |
| Walsh, 2018[84] | R L/S | bupropion 150mg | 20 | 20 | bw | bw | PILT | points won | 23.82 | 0.5 | 1 |
| Weis, 2012[85] | R L/S | L-dopa 100mg+35mg | 27 | 28 | bw | bw | appetitive instrumental conditioning | accuracy learning | 28.5 | 0.31 | 1 |
| Westbrook, 2024[48] | R L/S | methylphenidate 20mg | 95 | 97 | w† | bw | PILT WM | late accuracy | NA | NA | 1 |
| Addicott, 2021[72] | R S | methylphenidate 40mg | 22 | 22 | w | w | restless bandit | inverse temperature | 27.7 | 0.57 | 1 |
| Bernacer, 2013[2] | R S | methamphetamine 0.3mg/kg | 17 | 17 | w† | bw | PILT | temperature | 25.3 | 0.41 | 1 |
| Cremer, 2023[75] | R S | amisulpride 400mg | 23 | 22 | bw | bw | foraging | inverse temperature | NA | 0.47 | 1 |
| Eisenegger, 2014[13] | R S | sulpiride 800mg | 41 | 35 | bw | bw | PILT | temperature | NA | 0 | 1 |
| Admon, 2017*[73] | P L/S | amisulpride 50mg | 20 | 18 | bw | bw | PST | avoid B | 25.9 | 0.77 | 1 |
| Bellebaum, 2017[1] | P L/S | modafinil 200mg | 16 | 20 | bw | bw | PST | avoid B | 24.85 | 0 | 1 |
| Bernacer, 2013[2] | P L/S | methamphetamine 0.3mg/kg | 17 | 17 | w† | bw | PILT | learning rate | 25.3 | 0.41 | 1 |
| Cox, 2015[7] | P L/S | Phe / Tyr depletion 100g | 10 | 10 | w | w | PST | avoid B/D | NA | NA | 1 |

| | | | | | | | | | | | |
|---|---|---|---|---|---|---|---|---|---|---|---|
| Eisenegger, 2014[13] | P L/S | sulpiride 800mg | 41 | 35 | bw | bw | PILT | learning rate | NA | 0 | 1 |
| Frank, 2006*[14] | P L/S | haloperidol 2mg | 19 | 20 | w† | bw | PST | avoid B | 21 | 0.54 | 1 |
| Halahakoon, 2024[79] | P L/S | pramipexole 1mg/day | 21 | 19 | bw | bw | PILT | accuracy change pre-post intervention | 23.5 | 0.5 | >1 |
| Jocham, 2011*[53] | P L/S | amisulpride 200mg | 16 | 16 | w | w | PST | p(correct) lose-lose | 26.13 | 0 | 1 |
| Jocham, 2014[19] | P L/S | amisulpride 400mg | 22 | 22 | w | w | PILT | final accuracy | NA | 0 | 1 |
| Kelm, 2015[86] | P L/S | Phe / Tyr depletion 68.7g | 16 | 16 | w | w | PAL | passive avoidance errors | NA | 0.31 | 1 |
| Murray, 2019[83] | P L/S | amisulpride 400mg | 18 | 18 | w† | bw | PILT | accuracy | 32.1 | 0.17 | 1 |
| Pessiglione, 2006[26] | P L/S | L-dopa 100mg+25mg | 13 | 13 | bw | bw | PILT | accuracy | NA | NA | 1 |
| Walsh, 2018[84] | P L/S | bupropion 150mg | 20 | 20 | bw | bw | PILT | points lost | 23.82 | 0.5 | 1 |
| Bernacer, 2013[2] | P S | methamphetamine 0.3mg/kg | 17 | 17 | w† | bw | PILT | temperature | 25.3 | 0.41 | 1 |
| Eisenegger, 2014[13] | P S | sulpiride 800mg | 41 | 35 | bw | bw | PILT | temperature | NA | 0 | 1 |
| Guitart-Masip, 2012[16] | P+ | L-dopa 150mg+37.5mg | 16 | 20 | bw | bw | go/nogo | nogo to win | 23.15 | 0.34 | 1 |
| Guitart-Masip, 2014[17] | P+ | L-dopa 150mg+37.5mg | 30 | 29 | bw | bw | go/nogo | nogo to win | NA | NA | 1 |
| Hebart, 2015[18] | P+ | Phe / Tyr depletion | 35 | 34 | bw | bw | PIT | appetitive PIT | NA | NA | 1 |
| Scholz, 2022[32] | P+ | tolcapone 200mg | 35 | 35 | w† | bw | go/nogo | nogo to win | 31.3 | 0.26 | 1 |
| Weber, 2016[47] | P+ | amisulpride 400mg | 41 | 40 | bw | bw | PIT | button press rew-unrew CS | 21.8 | NA | 1 |
| Guitart-Masip, 2012[16] | P- | L-dopa 150mg+37.5mg | 16 | 20 | bw | bw | go/nogo | go to avoid | 23.15 | 0.34 | 1 |
| Guitart-Masip, 2014[17] | P- | L-dopa 150mg+37.5mg | 30 | 29 | bw | bw | go/nogo | go to avoid | NA | NA | 1 |
| Hebart, 2015[18] | P- | Phe / Tyr depletion | 35 | 34 | bw | bw | PIT | aversive PIT | NA | NA | 1 |
| Scholz, 2022[32] | P- | tolcapone 200mg | 35 | 35 | w† | bw | go/nogo | go to avoid | 31.3 | 0.26 | 1 |
| Daood, 2022[87] | RD | methylphenidate 20mg | 57 | 57 | w | w | DD | p(choose now) | 26.42 | 0.6 | 1 |
| Erfanian Abdoust, 2024*[88] | RD | haloperidol 2mg | 62 | 62 | w | w | DD | p(high cost option) | 22.79 | 0.52 | 1 |
| Kayser, 2012[89] | RD | tolcapone 200mg | 23 | 23 | w | w | DD | p(choose now) | NA | 0.56 | 1 |
| Kelm, 2013[90] | RD | Phe / Tyr depletion 68.7g | 15 | 15 | w | w | DD | p(choose now) | NA | 0 | 1 |
| Pine, 2010[65] | RD | L-dopa 150mg | 13 | 13 | w | w | DD | k parameter | 21 | 0.57 | 1 |
| Smith, 2024[91] | RD | L-dopa 150mg | 76 | 76 | w† | bw | DD | Larger-but-later choices | 28.26 | 0.42 | 1 |
| Soder, 2021[34] | RD | d-amphetamine 20mg | 28 | 28 | w† | bw | physical effort | p(hard task) | 24.07 | 0.5 | 1 |
| Wagner, 2020*[92] | RD | haloperidol 2mg | 23 | 26 | bw | bw | DD | p(delay) | 23.85 | 0.74 | 1 |
| Wardle, 2011[37] | RD | d-amphetamine 20mg | 17 | 17 | w† | bw | physical effort | p(hard task) | 22.7 | 0.65 | 1 |
| Weber, 2016[47] | RD | amisulpride 400mg | 40 | 40 | bw | bw | DD | p(choose now) | NA | NA | 1 |
| Westbrook, 2020[38] | RD | methylphenidate 20mg | 46 | 46 | w | w | cognitive effort | subjective value(hard task) | NA | NA | 1 |
| de Wit, 2002[93] | RD | d-amphetamine 20mg | 36 | 36 | w† | bw | DD | k parameter | 24 | 0.5 | 1 |
| Abler, 2007[45] | RV | olanzapine 5mg | 8 | 8 | w | w | MID | RT high reward | 31.3 | 0.5 | 1 |

| | | | | | | | | | | | |
|---|---|---|---|---|---|---|---|---|---|---|---|
| Au-Yeung, 2024[94] | RV | pramipexole 1mg/day | 19 | 18 | bw | bw | eye-tracking vigor | Velocity residuals | 23.59 | 0.48 | >1 |
| Beierholm, 2013[95] | RV | L-dopa 150mg+37.5mg | 30 | 30 | bw | bw | odd-ball discrimination | average reward rate on RTs | 24.15 | 0.43 | 1 |
| Graf, 2018[96] | RV | amisulpride 200mg/day | 17 | 17 | w | w | MID | RT 100-0 | 23.8 | 0 | >1 |
| Grimm, 2021[97] | RV | L-dopa 100mg+25mg | 45 | 45 | w | w | MID | RT high-neutral reward | 22.81 | 0.51 | 1 |
| Ikeda, 2019[98] | RV | bupropion 150mg | 15 | 15 | w | w | MID | RT high-neutral reward | 31.7 | 0.53 | 1 |
| Mehta, 2001[99] | RV | bromocriptine 1.25mg | 18 | 18 | w | w | Reward-responsivity | No. cards sorted in sort 3 - No. cards sorted in sorts 2/4 | NA | 0 | 1 |
| Mehta, 2005[100] | RV | Phe / Tyr depletion | 14 | 14 | w | w | Reward-responsivity | No. extra cards correctly sorted | 33.9 | 0.14 | 1 |
| Michely, 2020[46] | RV | L-dopa 150mg | 20 | 20 | w | w | dynamic effort | grip force high reward | 25.6 | 0.45 | 1 |
| Clatworthy, 2009[101] | MB/F | methylphenidate 60mg | 9 | 9 | w | w | reversal | reversal learning | NA | 0 | 1 |
| Deserno, 2021[10] | MB/F | L-dopa 150mg+37.5mg | 62 | 62 | w | w | model-based bandit | model-based choice | NA | NA | 1 |
| Ersche, 2011[102] | MB/F | amisulpride 400mg | 18 | 18 | w | w | probabilistic reversal | perseverative errors | 32.7 | 0.2 | 1 |
| Janssen, 2015[103] | MB/F | sulpiride 400mg | 22 | 22 | w | w | observational reversal | reversal errors | 32.2 | 0 | 1 |
| Mikus, 2022[24] | MB/F | amisulpride 400mg | 38 | 35 | bw | bw | Two-step task | omega parameter | NA | NA | 1 |
| Robinson, 2010[29] | MB/F | Phe / Tyr depletion | 27 | 27 | w | w | observational reversal | reversal errors | NA | 0.48 | 1 |
| Vo, 2018[104] | MB/F | L-dopa 100mg+25mg | 13 | 13 | bw | bw | probabilistic reversal | reversal errors | NA | NA | 1 |
| Wunderlich, 2012[40] | MB/F | L-dopa 150mg+37.5mg | 18 | 18 | w | w | Two-step task | omega parameter | 23.3 | 0 | 1 |
| van den Bosch, 2022[105] | MB/F | methylphenidate 20mg | 88 | 88 | w† | bw | observational reversal | accuracy reversal | NA | NA | 1 |
| van der Schaaf, 2013[106] | MB/F | methylphenidate 20mg | 19 | 19 | w | w | observational reversal | accuracy reversal | 20.9 | 0.52 | 1 |
| van der Schaaf, 2014[107] | MB/F | sulpiride 400mg | 23 | 23 | w | w | observational reversal | accuracy reversal | NA | 0.5 | 1 |
| Burke, 2018[66] | RA | amisulpride 400mg | 45 | 48 | bw | bw | gambling | utility curvature | NA | NA | 1 |
| Campbell-Meiklejohn, 2011*[108] | RA | pramipexole 0.176µg | 15 | 15 | bw | bw | gambling | p(gamble) | 24.9 | 0.53 | 1 |
| Hirschbichler, 2022[109] | RA | L-dopa 100mg | 30 | 30 | w | w | gambling | p(gamble) | 31.67 | 0.53 | 1 |
| Norbury, 2013[110] | RA | cabergoline 1.5mg | 20 | 20 | w | w | gambling | p(gamble) | 26.7 | NA | 1 |
| Riba, 2008*[111] | RA | pramipexole 0.5mg | 15 | 15 | w | w | gambling | p(risky choice) | 24.4 | NA | 1 |
| Rigoli, 2016[112] | RA | L-dopa 150mg+37.5mg | 32 | 32 | w | w | gambling | p(gamble) | 23.4 | 0.5 | 1 |
| Rutledge, 2015[113] | RA | L-dopa 150mg+37.5mg | 30 | 30 | w | w | gambling | p(gamble) gain | 23.4 | 0.63 | 1 |

**eTable 5: Characteristics of all dopamine studies.** Asterisks indicate studies that used a single low dose of an agonist or antagonists. These effects were interpreted contrary to their typical activity profile (e.g., a low-dose agonist acting as an antagonist), as suggested by the literature. Daggers indicate studies that were originally within-subject studies (in column 'Original design') but we could only calculate a between-subjects Cohen's *d* (indicated in column 'Final design'). In drug regimen, 1 indicates a single drug administration and >1 indicates

multiple drug administrations. Abbreviations: R L/S: reward learning/sensitivity; R S: reward sensitivity; P L/S: punishment learning/sensitivity; P S: punishment sensitivity; P+: appetitive Pavlovian; P-: aversive Pavlovian; RD: reward discounting; RV: reward vigor: MB/F: model-based learning/flexibility; RA: risk attitude; w: within-subject studies; bw: between-subject studies; PST: probabilistic selection task; PILT: probabilistic instrumental learning task; WM: working memory; PAL: passive avoidance learning; PIT: Pavlovian instrumental task; DD: delay discounting; MID: monetary incentive delay.

| Study | Category | Drug, Dose | N drug | N placebo | Original design | Final design | Task | Outcome | Study Age | Study Female Proportion | Drug regimen |
|---|---|---|---|---|---|---|---|---|---|---|---|
| Colwell, 2024[114] | R L/S | fenfluramine 15mg b.i.d. | 26 | 27 | bw | bw | PILT | learning rate | 20.17 | 0.6 | >1 |
| Cools, 2008[6] | R L/S | TD | 12 | 12 | w | w | observational reversal | non-reversal errors | 22.4 | 0.67 | 1 |
| Finger, 2007[115] | R L/S | TD | 16 | 19 | bw | bw | PAL | omission errors | 27.94 | 0.52 | 1 |
| Gilger, 2024[116] | R L/S | TD | 98 | 98 | w† | bw | Two-step | main effect reward | 32.2 | 0.38 | 1 |
| Jepma, 2010[117] | R L/S | citalopram 30mg | 16 | 17 | bw | bw | restless bandit | inverse temperature | 21.55 | 0.49 | 1 |
| Luo, 2024[67] | R L/S | escitalopram 20mg | 32 | 33 | bw | bw | probabilistic reversal | learning rate | 26 | 0.49 | 1 |
| Michely, 2022[118] | R L/S | citalopram 20mg/day | 33 | 33 | bw | bw | gambling card | learning rate | 24.65 | 0.39 | >1 |
| Robinson, 2012[28] | R L/S | TD | 21 | 20 | bw | bw | observational reversal | non-reversal errors | 27.6 | 1 | 1 |
| Seymour, 2012[33] | R L/S | TD | 15 | 15 | bw | bw | restless bandit | sensitivity | NA | NA | 1 |
| Colwell, 2024[114] | R S | fenfluramine 15mg b.i.d. | 26 | 27 | bw | bw | PILT | outcome sensitivity | 20.17 | 0.6 | >1 |
| Colwell, 2024[114] | P L/S | fenfluramine 15mg b.i.d. | 26 | 27 | bw | bw | PILT | learning rate | 20.17 | 0.6 | >1 |
| Luo, 2024[67] | P L/S | escitalopram 20mg | 32 | 33 | bw | bw | probabilistic reversal | learning rate | 26 | 0.49 | 1 |
| Michely, 2022[118] | P L/S | citalopram 20mg/day | 33 | 33 | bw | bw | gambling card | learning rate | 24.65 | 0.39 | >1 |
| Seymour, 2012[33] | P L/S | TD | 15 | 15 | bw | bw | restless bandit | sensitivity | NA | NA | 1 |
| Colwell, 2024[114] | P S | fenfluramine 15mg b.i.d. | 26 | 27 | bw | bw | PILT | outcome sensitivity | 20.17 | 0.6 | >1 |
| Geurts, 2013[15] | P+ | TD | 45 | 45 | w | w | PIT | appetitive PIT | 23.8 | NA | 1 |
| Guitart-Masip, 2012[16] | P+ | citalopram 24mg oral | 16 | 20 | bw | bw | go/nogo | nogo to win | 23.3 | 0.49 | 1 |
| Guitart-Masip, 2014[17] | P+ | citalopram 30mg | 27 | 29 | bw | bw | go/nogo | nogo to win | NA | NA | 1 |
| Hebart, 2015[18] | P+ | TD | 34 | 34 | bw | bw | PIT | appetitive PIT | NA | NA | 1 |
| Cools, 2008[6] | P- | TD | 12 | 12 | w | w | observational reversal | non-reversal errors | 22.4 | 0.67 | 1 |
| Crockett, 2009[9] | P- | TD | 22 | 22 | w | w | go/nogo RT | punishment-induced slowing | NA | NA | 1 |
| Crockett, 2012[8] | P- | TD | 24 | 24 | w | w | go/nogo RT | punishment-induced slowing | NA | NA | 1 |
| Gaber, 2015[119] | P- | TD | 24 | 24 | w | w | go/nogo RT | punishment-induced slowing | 25.34 | NA | 1 |
| Geurts, 2013[15] | P- | TD | 45 | 45 | w | w | PIT | aversive PIT | 23.8 | NA | 1 |
| Guitart-Masip, 2012[16] | P- | citalopram 24mg oral | 16 | 20 | bw | bw | go/nogo | go to avoid | 23.3 | 0.49 | 1 |

| | | | | | | | | | | | |
|---|---|---|---|---|---|---|---|---|---|---|---|
| Guitart-Masip, 2014[17] | P- | citalopram 30mg | 27 | 29 | bw | bw | go/nogo | go to avoid | NA | NA | 1 |
| Hebart, 2015[18] | P- | TD | 34 | 34 | bw | bw | PIT | aversive PIT | NA | NA | 1 |
| Helmbold, 2015[120] | P- | TD | 18 | 18 | w | w | go/nogo RT | punishment-induced slowing | 24.22 | NA | 1 |
| Robinson, 2012[28] | P- | TD | 21 | 20 | bw | bw | observational reversal | non-reversal errors | 27.6 | 1 | 1 |
| Crean, 2002[121] | RD | TD | 20 | 20 | w† | bw | DD | k parameter | 21.5 | 0 | 1 |
| Schweighofer, 2008[122] | RD | TD | 20 | 20 | w | w | DD dynamic | discount factor gamma | NA | 0 | 1 |
| Tanaka, 2007[123] | RD | TD | 12 | 12 | w† | bw | multi-step DD | p(large reward choices) | NA | 0 | 1 |
| Worbe, 2014[124] | RD | TD | 22 | 22 | bw | bw | DD | k parameter | 28.97 | 0.61 | 1 |
| Beierholm, 2013[95] | RV | citalopram 24mg oral | 30 | 30 | bw | bw | odd-ball discrimination | average reward rate on RTs | 23.91 | 0.5 | 1 |
| Cools, 2005[125] | RV | TD | 10 | 12 | bw | bw | CRRT | RT 90-10 | 23.55 | 0 | 1 |
| Roiser, 2006[126] | RV | TD | 30 | 30 | w | w | CRRT | RT 90-10 | 26.75 | 0.43 | 1 |
| Steding, 2023[127] | RV | TD | 24 | 24 | w | w | physical effort | button press high-low | NA | 1 | 1 |
| Chamberlain, 2006[128] | MB/F | citalopram 30mg | 20 | 20 | bw | bw | reversal | perseverative errors | 25.82 | 0.5 | 1 |
| Cools, 2008[6] | MB/F | TD | 12 | 12 | w | w | observational reversal | switch errors | 22.4 | 0.67 | 1 |
| Evers, 2004[129] | MB/F | TD | 15 | 15 | w† | bw | probabilistic reversal | perseverative errors | 21.8 | 0.8 | 1 |
| Gilger, 2024[116] | MB/F | TD | 98 | 98 | w† | bw | Two-step | omega parameter | 32.2 | 0.38 | 1 |
| Kanen, 2020[130] | MB/F | TD | 30 | 32 | bw | bw | probabilistic reversal | perseveration | NA | 0.47 | 1 |
| Langley, 2023[131] | MB/F | escitalopram 20mg /day | 32 | 34 | bw | bw | Two-step task | Reward × Transition | 25 | 65.17 | >1 |
| Skandali, 2018[132] | MB/F | escitalopram 20mg | 31 | 33 | bw | bw | Probabilistic reversal | reversal errors | 26 | 0.49 | 1 |
| Thirkettle, 2019[133] | MB/F | Tryptophan 0.8g | 22 | 22 | bw | bw | probabilistic reversal | reversal errors | 21.1 | 0.64 | 1 |
| Worbe, 2015[134] | MB/F | TD | 18 | 18 | bw | bw | 3-stage instrumental learning | devalued choices | 26.33 | 0.56 | 1 |
| Worbe, 2016[39] | MB/F | TD | 22 | 22 | bw | bw | Two-step task | omega parameter | 29.14 | 0.52 | 1 |
| Anderson, 2003[135] | RA | TD | 15 | 13 | bw | bw | gambling | mean indifference odds against win | 23 | 0.39 | 1 |
| Campbell-Meiklejohn, 2011[108] | RA | TD | 17 | 17 | bw | bw | gambling | p(gamble) | 24.18 | 0.53 | 1 |
| Faulkner, 2017[136] | RA | TD | 26 | 26 | w | w | gambling | p(gamble) | 29.45 | 0.54 | 1 |
| Macoveanu, 2014[137] | RA | fluoxetine 20/40mg/day | 13 | 16 | bw | bw | gambling | p(risky choice) | NA | 0 | >1 |
| Murphy, 2009[138] | RA | l-tryptophan 3g/day | 15 | 15 | bw | bw | gambling | p(gamble) | 25.54 | 0.53 | >1 |
| Rogers, 2003[111] | RA | TD | 18 | 18 | bw | bw | gambling | p(gamble) | 23.8 | 0.5 | 1 |

**eTable 6: Characteristics of all serotonin studies.** Daggers indicate studies that were originally within-subject studies (in column ’Original design’) but we could only calculate a between-subjects Cohen’s *d* (indicated in column ’Final design’). In drug regimen, 1 indicates a single drug administration and >1 indicates multiple drug administrations. Abbreviations: R L/S: reward

learning/sensitivity; R S: reward sensitivity; P L/S: punishment learning/sensitivity; P S: punishment sensitivity; P+: appetitive Pavlovian; P-: aversive Pavlovian; RD: reward discounting; RV: reward vigor: MB/F: model-based learning/flexibility; RA: risk attitude; w: within-subject studies; bw: between-subject studies; PST: probabilistic selection task; PILT: probabilistic instrumental learning task; PAL: passive avoidance learning; PIT: Pavlovian instrumental task; DD: delay discounting; MID: monetary incentive delay; CRRT: cued-reinforcement reaction-time task.

### 4 eAppendix 1

#### 4.1 PRISMA 2020 Checklist

| Section and Topic | Item # | Checklist item | Location where item is reported |
|---|---|---|---|
| **TITLE** | | | |
| Title | 1 | Identify the report as a systematic review. | Title |
| **ABSTRACT** | | | |
| Abstract | 2 | See the PRISMA 2020 for Abstracts checklist. | Appendix 1: 4.2 PRISMA 2020 Abstract Checklist |
| **INTRODUCTION** | | | |
| Rationale | 3 | Describe the rationale for the review in the context of existing knowledge. | Introduction |
| Objectives | 4 | Provide an explicit statement of the objective(s) or question(s) the review addresses. | Introduction |
| **METHODS** | | | |
| Eligibility criteria | 5 | Specify the inclusion and exclusion criteria for the review and how studies were grouped for the syntheses. | Methods, Supplementary materials: 1.2 Paper selection, 1.3 Meta-analysis |
| Information sources | 6 | Specify all databases, registers, websites, organisations, reference lists and other sources searched or consulted to identify studies. Specify the date when each source was last searched or consulted. | Methods |
| Search strategy | 7 | Present the full search strategies for all databases, registers and websites, including any filters and limits used. | eTable 1 |
| Selection process | 8 | Specify the methods used to decide whether a study met the inclusion criteria of the review, including how many reviewers screened each record and each report retrieved, whether they worked independently, and if applicable, details of automation tools used in the process. | Supplementary methods: 1.2 Paper selection |
| Data collection process | 9 | Specify the methods used to collect data from reports, including how many reviewers collected data from each report, whether they worked independently, any processes for obtaining or confirming data from study investigators, and if applicable, details of automation tools used in the process. | Supplementary methods: 1.2 Paper selection, 1.3 Meta-analysis |
| Data items | 10a | List and define all outcomes for which data were sought. Specify whether all results that were compatible with each outcome domain in each study were sought (e.g. for all measures, time points, analyses), and if not, the methods used to decide which results to collect. | Methods, Supplementary methods: 1.2 Paper selection, 1.3 Meta-analysis |
| | 10b | List and define all other variables for which data were sought (e.g. participant and intervention characteristics, funding sources). Describe any assumptions made about any missing or unclear information. | eTable5-6 |
| Study risk of bias assessment | 11 | Specify the methods used to assess risk of bias in the included studies, including details of the tool(s) used, how many reviewers assessed each study and whether they worked independently, and if applicable, details of automation tools used in the process. | Supplementary methods: 1.3.5 Assessment of study quality |
| Effect measures | 12 | Specify for each outcome the effect measure(s) (e.g. risk ratio, mean difference) used in the synthesis or presentation of results. | Methods |

| Section and Topic | Item # | Checklist item | Location where item is reported |
|---|---|---|---|
| Synthesis methods | 13a | Describe the processes used to decide which studies were eligible for each synthesis (e.g. tabulating the study intervention characteristics and comparing against the planned groups for each synthesis (item #5)). | Methods, Supplementary methods: 1.2 Paper selection, 1.3 Meta-analysis |
| | 13b | Describe any methods required to prepare the data for presentation or synthesis, such as handling of missing summary statistics, or data conversions. | Supplementary methods: 1.4 Effect size calculations |
| | 13c | Describe any methods used to tabulate or visually display results of individual studies and syntheses. | Figure 2-5, eFigure 3-13 |
| | 13d | Describe any methods used to synthesize results and provide a rationale for the choice(s). If meta-analysis was performed, describe the model(s), method(s) to identify the presence and extent of statistical heterogeneity, and software package(s) used. | Methods, Supplementary methods: 1.3 Meta-analysis |
| | 13e | Describe any methods used to explore possible causes of heterogeneity among study results (e.g. subgroup analysis, meta-regression). | Supplementary methods: 1.3 Meta-analysis |
| | 13f | Describe any sensitivity analyses conducted to assess robustness of the synthesized results. | Methods, Supplementary methods: 1.3 Meta-analysis |
| Reporting bias assessment | 14 | Describe any methods used to assess risk of bias due to missing results in a synthesis (arising from reporting biases). | Supplementary methods: 1.3 Meta-analysis |
| Certainty assessment | 15 | Describe any methods used to assess certainty (or confidence) in the body of evidence for an outcome. | Supplementary methods: 1.3.5 Assessment of study quality |
| **RESULTS** | | | |
| Study selection | 16a | Describe the results of the search and selection process, from the number of records identified in the search to the number of studies included in the review, ideally using a flow diagram. | Figure 1 |
| | 16b | Cite studies that might appear to meet the inclusion criteria, but which were excluded, and explain why they were excluded. | Supplementary methods: 1.2 Paper selection, Figure 1 |
| Study characteristics | 17 | Cite each included study and present its characteristics. | Figure 2-5, eFigure 3-13, eTable 5-6 |
| Risk of bias in studies | 18 | Present assessments of risk of bias for each included study. | eFigure 2 |
| Results of individual studies | 19 | For all outcomes, present, for each study: (a) summary statistics for each group (where appropriate) and (b) an effect estimate and its precision (e.g. confidence/credible interval), ideally using structured tables or plots. | Figure 2-5, eFigure 3-13 |
| Results of syntheses | 20a | For each synthesis, briefly summarise the characteristics and risk of bias among contributing studies. | Results |
| | 20b | Present results of all statistical syntheses conducted. If meta-analysis was done, present for each the summary estimate and its precision (e.g. confidence/credible interval) and measures of statistical heterogeneity. If comparing groups, describe the direction of the effect. | Results, Figure 2-5, eFigure 3-13, Supplementary results |

| Section and Topic | Item # | Checklist item | Location where item is reported |
|---|---|---|---|
| | 20c | Present results of all investigations of possible causes of heterogeneity among study results. | Results, Supplementary results: 2.3 Dopamine heterogeneity, 2.5 serotonin heterogeneity, 2.6 Moderator analyses, 2.7 Publication bias |
| | 20d | Present results of all sensitivity analyses conducted to assess the robustness of the synthesized results. | Supplementary results: 2.2 Additional dopamine results, 2.4 Additional serotonin results, 2.8 Additional supplementary analyses |
| Reporting biases | 21 | Present assessments of risk of bias due to missing results (arising from reporting biases) for each synthesis assessed. | Results, Supplementary methods: 2.7 Publication bias |
| Certainty of evidence | 22 | Present assessments of certainty (or confidence) in the body of evidence for each outcome assessed. | Results, Figures 2-5, eFigure 3-13 |
| **DISCUSSION** | | | |
| Discussion | 23a | Provide a general interpretation of the results in the context of other evidence. | Discussion |
| | 23b | Discuss any limitations of the evidence included in the review. | Discussion |
| | 23c | Discuss any limitations of the review processes used. | Discussion |
| | 23d | Discuss implications of the results for practice, policy, and future research. | Discussion |
| **OTHER INFORMATION** | | | |
| Registration and protocol | 24a | Provide registration information for the review, including register name and registration number, or state that the review was not registered. | Methods, Additional information |
| | 24b | Indicate where the review protocol can be accessed, or state that a protocol was not prepared. | Additional information |
| | 24c | Describe and explain any amendments to information provided at registration or in the protocol. | In the preregistered protocol |
| Support | 25 | Describe sources of financial or non-financial support for the review, and the role of the funders or sponsors in the review. | Financial disclosures |
| Competing interests | 26 | Declare any competing interests of review authors. | Acknowledgments |
| Availability of data, code and other materials | 27 | Report which of the following are publicly available and where they can be found: template data collection forms; data extracted from included studies; data used for all analyses; analytic code; any other materials used in the review. | Additional information |

#### 4.2 PRISMA 2020 Abstract Checklist

| Section and Topic | Item # | Checklist item | Reported (Yes/No) |
|---|---|---|---|
| **TITLE** | | | |
| Title | 1 | Identify the report as a systematic review. | Yes |
| **BACKGROUND** | | | |
| Objectives | 2 | Provide an explicit statement of the main objective(s) or question(s) the review addresses. | Yes |
| **METHODS** | | | |
| Eligibility criteria | 3 | Specify the inclusion and exclusion criteria for the review. | Yes |
| Information sources | 4 | Specify the information sources (e.g. databases, registers) used to identify studies and the date when each was last searched. | Yes |
| Risk of bias | 5 | Specify the methods used to assess risk of bias in the included studies. | Yes |
| Synthesis of results | 6 | Specify the methods used to present and synthesise results. | Yes |
| **RESULTS** | | | |
| Included studies | 7 | Give the total number of included studies and participants and summarise relevant characteristics of studies. | Yes |
| Synthesis of results | 8 | Present results for main outcomes, preferably indicating the number of included studies and participants for each. If meta-analysis was done, report the summary estimate and confidence/credible interval. If comparing groups, indicate the direction of the effect (i.e. which group is favoured). | Yes |
| **DISCUSSION** | | | |
| Limitations of evidence | 9 | Provide a brief summary of the limitations of the evidence included in the review (e.g. study risk of bias, inconsistency and imprecision). | No |
| Interpretation | 10 | Provide a general interpretation of the results and important implications. | Yes |
| **OTHER** | | | |
| Funding | 11 | Specify the primary source of funding for the review. | No |
| Registration | 12 | Provide the register name and registration number. | No |
